# Arrayed single-gene perturbations identify drivers of human anterior neural tube closure

**DOI:** 10.1101/2025.07.21.665862

**Authors:** Roya E. Huang, Giridhar M. Anand, Heitor C. Megale, Jason Chen, Chudi Abraham-Igwe, Sharad Ramanathan

## Abstract

Genetic studies of human embryonic morphogenesis are constrained by ethical and practical challenges, restricting insights into developmental mechanisms and disorders. Human pluripotent stem cell (hPSC)–derived organoids provide a powerful alternative for the study of embryonic morphogenesis. However, screening for genetic drivers of morphogenesis *in vitro* has been infeasible due to organoid variability and the high costs of performing scaled tissue-wide single-gene perturbations. By overcoming both these limitations, we developed a platform that integrates reproducible organoid morphogenesis with uniform single-gene perturbations, enabling high-throughput arrayed CRISPR interference (CRISPRi) screening in hPSC-derived organoids. To demonstrate the power of this platform, we screened 77 transcription factors in an organoid model of anterior neurulation to identify *ZIC2*, *SOX11*, and *ZNF521* as essential regulators of neural tube closure. We discovered that *ZIC2* and *SOX11* are required for closure, while *ZNF521* prevents ectopic closure points. Single-cell transcriptomic analysis of perturbed organoids revealed co-regulated gene targets of *ZIC2* and *SOX11* and an opposing role for *ZNF521*, suggesting that these transcription factors jointly govern a gene regulatory program driving neural tube closure in the anterior forebrain region. Our single-gene perturbation platform enables high-throughput genetic screening of *in vitro* models of human embryonic morphogenesis.

## Introduction

Pioneering genetic screens in model organisms—including notable work in *Drosophila* ^1^ and zebrafish ^2^—demonstrate that key regulators of embryonic morphology and tissue patterning are revealed by organism-wide gene perturbations. Performing analogous screens in human embryos is prohibited for ethical reasons, and screens in mammalian model organisms present significant challenges: generating clonal knockout lines in mice is expensive ^3–5^ and direct translation to human biology is complicated by species-specific differences in gene regulation, developmental timing, and tissue organization ^6^.

Human pluripotent stem cell (hPSC)–derived organoids offer an attractive alternative for modeling human embryonic morphogenesis *in vitro*. Advances in three-dimensional differentiation have enabled the generation of increasingly complex organoid models that recapitulate the development of different human tissues ^7–13^. The growing reproducibility of these models following the incorporation of bioengineering methods ^14–18^ makes them ideally suited for genetic perturbation screens. Despite progress, it remains challenging to interrogate the effects of genes on morphogenesis at scale. High-throughput screens to understand gene function in organoids have relied on CRISPR/Cas9-based approaches in which organoids are transduced with a pooled lentiviral library, with each virus carrying guide RNAs targeting a distinct gene. The approach produces mosaic organoids where multiple genes are knocked down in distinct cell sub-populations within the same organoid ^19–21^. This degree of mosaicism makes it impossible to study tissue-wide morphogenesis, which occurs via the coordination of genes acting across many cells. Pooled screens are limited to single-cell genomic, epigenomic, or transcriptomic profiling as a readout of gene function, in contrast to morphological phenotypes assessed by classical screens in developmental biology.

The study of morphogenesis at scale *in vitro* requires the ability to knock out or knock down genes individually throughout entire organoids. This typically requires the generation of knockout or knockdown cell lines through clonal analysis, a process that is prohibitive for high-throughput applications due to the cost and labor involved in isolating and expanding correctly edited clones ^22–24^. The ability to achieve homogeneous gene knockdown across a cell population in a non-clonal manner—using technologies such as CRISPR interference (CRISPRi)—could enable high-throughput morphological screens ^25^. Lentiviruses carrying CRISPRi guide RNAs offer a means to perform such non-clonal knockdowns, and if delivered with high efficiency, can eliminate the need for clonal purification and expansion. However, this strategy faces several challenges: current protocols to generate high-titer lentivirus relies on laborious and time-consuming concentration steps which are not amenable to high-throughput workflows ^26^, and it is difficult to achieve high-efficiency lentiviral transduction without associated cell death ^27^. Overcoming these critical challenges would enable genetic perturbations at a scale not currently feasible in organoids or mammalian model systems, allowing systematic interrogation of the genes that drive human embryonic morphogenesis.

Here, by developing a method to generate and apply high-titer lentivirus directly to stem cells in parallelized small volumes, we established a high-throughput arrayed CRISPRi platform for uniform knockdown of individual genes across whole hPSC-derived organoids. To demonstrate the utility of this platform, we applied it to study anterior neurulation, a critical morphogenetic event in brain formation in which the flat anterior neural plate bends and fuses to form a closed neural tube ^28,29^. Defects in anterior neural tube closure cause anencephaly, which is embryonic lethal. This lethality limits genetic linkage and association studies ^30^ and makes anterior neurulation of critical importance to study *in vitro*. To systematically determine genes required for anterior neurulation, we established a robust *in vitro* hPSC-derived anterior neural tube organoid model and used our CRISPRi platform to screen 77 individual candidate genes encoding transcription factors for their contribution to neural tube morphogenesis. We found that *ZIC2* and *SOX11* are required for anterior neural tube closure and co-regulate a gene expression program in opposition to *ZNF521*, which is required to prevent ectopic closure points. These findings show the impact of scalable single-gene perturbations in identifying genes driving morphogenetic processes *in vitro* and have direct implications for understanding the genetic causes of congenital malformations.

## Results

### A reproducible *in vitro* model of human anterior neurulation

We previously discovered that when spatially arranged human pluripotent stem cell (hPSC)-derived epiblast-like cysts are exposed to posteriorizing signals, they break symmetry and undergo patterning ^14,18^. Building on this approach, we established an organoid model of human anterior neurulation (Figure 1A). Briefly, we seeded hPSCs onto a glass coverslip containing hexagonally arranged 250-micron-diameter micropatterns of extracellular matrix (Matrigel), including a central micropattern (Figure S1A), with 100-micron edge-to-edge spacing. After allowing hPSCs to form two-dimensional epithelial colonies on micropatterns, we embedded them in concentrated Matrigel, prompting hPSC colonies to fold into three-dimensional epithelial cysts with apicobasal polarity and single lumens. To induce anterior neural and surface ectoderm fates, we inhibited BMP, TGFβ, and WNT signaling for two days, then added recombinant human BMP4 for four days with continued TGFβ and WNT inhibition (Figure S1B-S1D, Methods) ^16,26,31^. Under these signaling conditions, the spatially arranged cysts differentiated into organoids with inward-facing neural ectoderm (NCAD+) and outward-facing surface ectoderm (ECAD+) (Figure 1B-C). Over the course of four days of BMP4 exposure, differentiating cysts displayed hallmarks of anterior primary neurulation, including thickening and elevation of the neural plate (Day 2), basal apposition of the neural and surface ectoderm (Day 3), and fusion of the neural folds at the midline to form a closed neural tube with overlying surface ectoderm (Day 4) (Figure 1B, 1C, S1E). In replicate experiments, all organoids displayed a closed neural lumen as assayed by a closed ring of apical NCAD on Day 4 (Figure 1D and S1F, n=24). Furthermore, quantification of features including organoid size, lumen size, and ECAD/NCAD area ratio showed that our *in vitro* organoids were reproducible with respect to tissue patterning and morphogenesis (Figure 1C, 1E, S1E, N=4 biological replicates, n=24 organoids).

**Figure 1.**
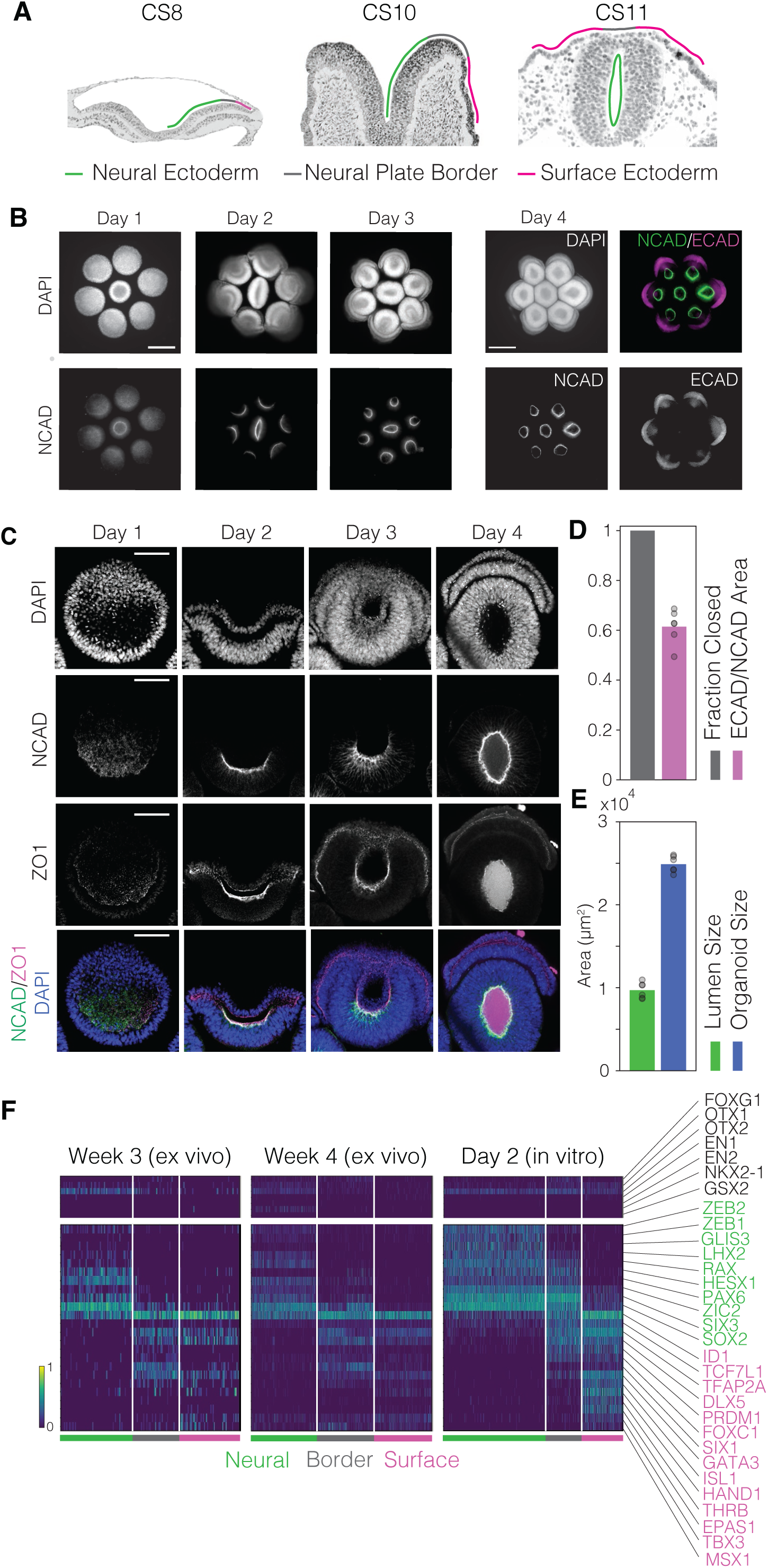
**An *in vitro* model of human anterior neurulation.** (A) Micrographs of human embryo sections from the Virtual Human Embryo project showing sequential neural plate bending, neural fold elevation, and neural tube closure from Carnegie Stages (CS) 8, 10, and 11. Tissues are outlined according to epithelial thickness, denoting the approximate location of neural ectoderm (green), neural plate border (gray), and surface ectoderm (magenta). Scale bar: 300 µm. (B) Top-down epifluorescence images of seven three-dimensional, hexagonally arranged neural tube organoids 1, 2, and 3 days after BMP4 addition, stained for nuclear marker DAPI and neural ectoderm marker N-cadherin (NCAD). Rightmost set of four images shows the organoids at day 4 of differentiation, stained for DAPI, NCAD, and surface ectoderm marker E-cadherin (ECAD). The six outer organoids show radial patterning, with inward-facing neural ectoderm (NCAD+, green) and outward-facing surface ectoderm (ECAD+, magenta). Scale bar: 300 µm. (C) Top-down confocal images of a representative single outer organoid after exposure to BMP4 for 1, 2, 3, and 4 days, fixed and stained for DAPI, NCAD, epithelial tight junction marker ZO-1. A neural plate is distinguishable by Day 2, which elevates and folds from a “U” shape to a “C” shape by Day 3 and finally fuses on Day 4 to enclose a single lumen (“O” shape). Extraneous ZO-1 staining in the lumen on Day 4 is likely due to dead cells that have been shed into the lumen during and after folding (see Supplementary Videos 1 and 2). Scale bar: 100 µm. (D) Quantification of neural tube closure in Day 4 organoids (left) and relative proportion of ECAD+ surface ectoderm to NCAD+ neural ectoderm (right) as measured by the projected area (see Methods). 100% of the outer organoids across biological replicates (N=4 replicates, n=24 organoids) exhibit closure based on observation of a continuous ring of NCAD (Figure S1F). The ratio of the surface to neural ectoderm has a coefficient of variation (CV) of 0.01. E) Quantification of lumen (left) and organoid size (right) across Day 4 organoids (n=6), based on the projected area enclosed by NCAD and DAPI staining (Methods). Lumen and organoid size have a CV of 0.09 and 0.04, respectively. F) Hierarchical clustering in the space of genes identified by sparse multimodal decomposition (Methods) of single-cell RNA-sequencing data from Week 3 human embryos, Week 4 human embryos, and organoids on Day 2 after BMP4 addition, showing the existence of neural ectoderm (green), neural plate border (gray) and surface ectoderm (magenta) cell types *in vitro* and *ex vivo* marked by similar gene expression patterns. Distinct sets of transcription factors are enriched in neural and surface ectoderm, highlighted in green and magenta respectively, and a subset of these are co-expressed in border ectoderm. Neural ectoderm cells express high levels of forebrain-associated transcription factors (OTX1, OTX2, LHX2, SIX3), and low levels of midbrain/hindbrain- (EN1, EN2) or ventral forebrain- (NKX2-1, GSX2) associated transcription factors. Gene expression (color bar, bottom left) is normalized to median transcript count, followed by log- and min-max normalization.

To characterize the cell types in our organoid model at the onset of neurulation, we performed single-cell RNA-sequencing (scRNA-seq) at Day 2 of BMP4 exposure, when the neural plate begins to elevate. To overcome limitations of existing computational methods in identifying cell types from scRNA-seq data, we applied our previously validated unsupervised dimensionality reduction method, sparse multimodal decomposition (SMD) ^32^. Using SMD, we identified a set of bimodally expressed genes that separated the cells into three cell types distributed along the mediolateral axis within the ectoderm: neural ectoderm, neural plate border, and surface ectoderm (Methods, Figure 1F, left). The three cell types showed a graded expression of *SOX2* in the medial-to-lateral direction: highest in the neural ectoderm, intermediate in the neural plate border, and lowest in the surface ectoderm, consistent with studies in model organisms ^33^. Neural ectoderm cells expressed forebrain-associated transcription factors, including *LHX2, HESX1, PAX6*, *SIX3*, *ZEB1* and *ZEB2* ^34–40^. Surface ectoderm cells expressed canonical transcription factors including *TFAP2A* and *GATA3* ^34,41^. Meanwhile, the neural plate border population co-expressed neural ectoderm transcription factors *PAX6* and *SIX3* as well as surface ectoderm transcription factors *TFAP2A* and *GATA3*, along with *DLX5* and *SIX1*, which mark cranial placodes arising at the neural plate border ^42,43^. Notably, all ectodermal cell types expressed *OTX1* and *OTX2* but not *EN1* or *EN2*, indicating forebrain rather than midbrain/hindbrain identities ^44^. Additionally, ventral markers *NKX2*-*1* and *GSX2* were absent, suggesting a dorsal identity ^45^. We did not observe neural crest cells, marked by transcription factors *SOX10* and *FOXD3* ^46,47^. This is consistent with neural tube closure in the human embryo at the anterior forebrain level, where no neural crest cells are present ^48^ . To compare our findings with *ex vivo* gene expression patterns, we applied the same approach to published scRNA-seq data from 3 to 4-week-old human embryos at the neurula stage^49^. Using the same SMD approach, we identified the same three ectodermal cell types *ex vivo*, with gene expression patterns that matched those observed in our organoids (Figure 1F). We conclude that our organoid model consists of cell types involved in human anterior neurulation. Cells from embryos showed higher expression of hemoglobin subunits and other hypoxia-related genes, which may indicate hypoxic stress from sample handling or reflect differences in the metabolic environment *ex vivo* (Fig. S1G). ^50^

### Identifying transcription factor candidates for regulation of anterior neurulation

Using our organoid model, we aimed to identify transcription factors driving anterior neurulation through a knockdown screen. Classical studies that physically disrupt the ectoderm during the neurula stages in vertebrates suggest that the neural ectoderm drives folding and closure of the neural plate (through elongation, tractoring, and/or bending), while the surface ectoderm is relatively passive ^51–53^. Therefore, we sought to identify transcription factors expressed within the neural ectoderm that drive this morphogenetic process. We ordered ectoderm cells along a pseudo-mediolateral coordinate using a diffusion map on scRNA-seq data from Day 2 (Figure 2A, left). We identified transcription factors expressed above a mean expression threshold across all cells and ranked them by the correlation of their expression with the mediolateral coordinate. This analysis revealed 20 transcription factors with highly specific expression in the neural plate (Figure 2A, right; Figure S2A, S2G, S2H). Thus, we selected these 20 transcription factors as candidates for a targeted knockdown screen.

**Figure 2.**
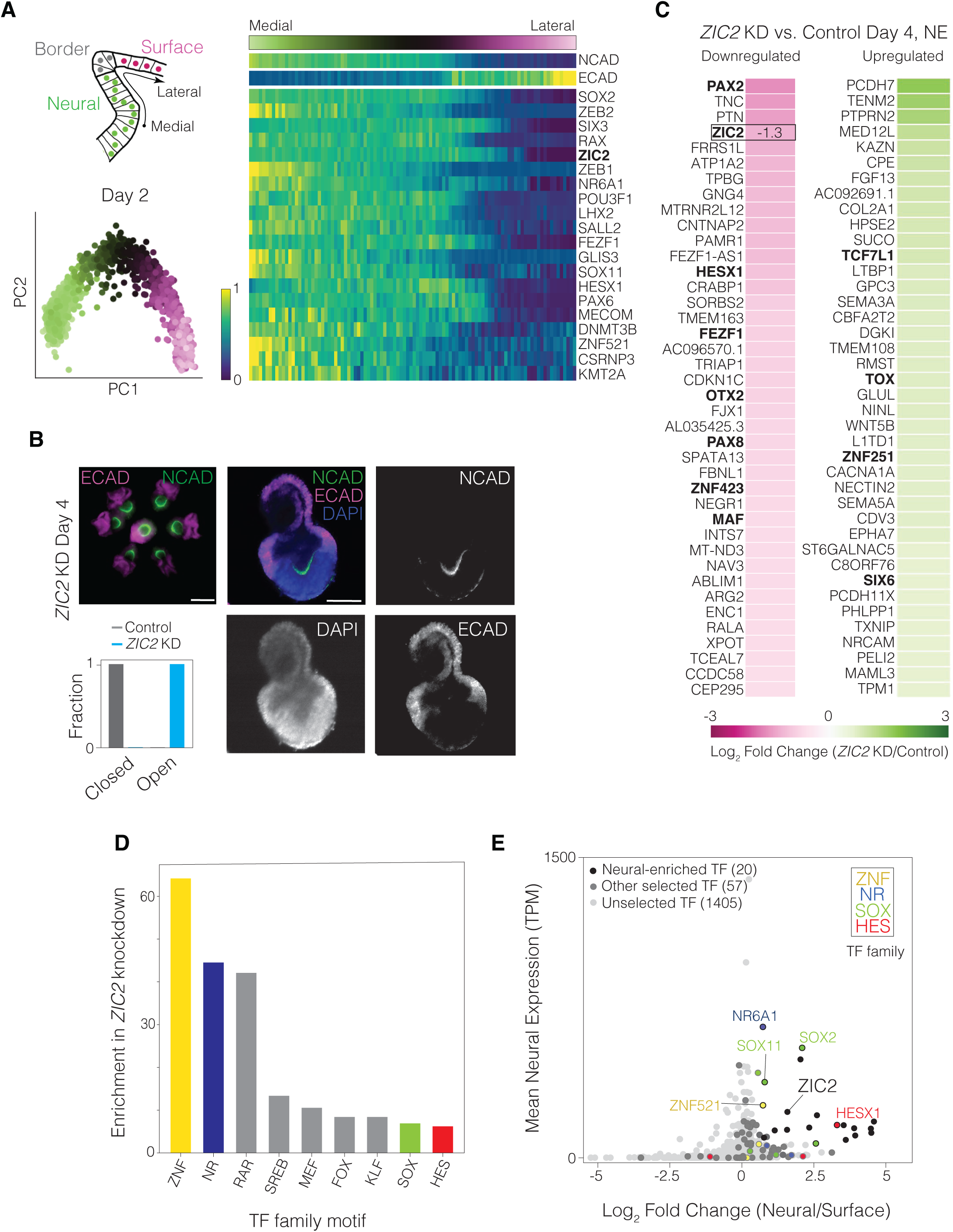
**Identifying transcription factor candidates regulating anterior neurulation.** A) Top left: Illustration denoting the mediolateral axis present in neural tube organoids, spanning the dorsal neural ectoderm, neural plate border, and surface ectoderm. Bottom left: Principal component analysis of single-cell RNA-sequencing (scRNA-seq) data from Day 2 organoids, colored by pseudo-spatial mediolateral coordinate. Right: Standardized mediolateral expression profiles of NCAD, ECAD, and 20 most medially expressed transcription factors from Day 2 organoids. Mediolateral coordinate is denoted by the green-magenta color bar (top) and expression is denoted by the blue-yellow color bar (bottom left). B) Day 4 organoids transduced with a dual-guide RNA targeting *ZIC2*. Top left: Epifluorescence merged-channel image of organoids on the pattern stained for NCAD and ECAD. Scale bar: 300 µm. Right: Confocal image of one organoid stained for NCAD, ECAD, and DAPI. Scale bar: 100 µm. Bottom left: Quantification of neural tube closure in Day 4 control and ZIC2 knockdown (KD) organoids, showing that 0% of the control and 100% of ZIC2 knockdown organoids possess an open neural plate. C) Top 40 up- and down-regulated genes in the neural ectoderm (NE) of Day 4 organoids upon *ZIC2* knockdown as compared to scramble control, based on log2(fold change) (color bar, bottom). Transcription factors highlighted in bold. D) Histogram of enrichment scores based on MEME SEA (Methods) of most overrepresented transcription factor family motifs in regulatory regions 10kb upstream and 100bp downstream of transcriptional start sites for the most down-regulated genes upon *ZIC2* knockdown, using a threshold of log2(mean fold change relative to scramble control) < −0.4. Families are color-coded if a transcription factor from that family is present in our candidate selection (Figure 2E). E) Plot of mean expression in neural ectoderm (transcripts per million, TPM) versus log2(fold change) in their expression between the neural and surface ectoderm in Day 2 organoids for all transcription factors (n=1482). Top 20 most medially expressed transcription factors (black or black outline) and additional candidates (gray, n=58) selected for knockdown (n=78, total). Data points corresponding to members of TF family with enrichment of binding sites near ZIC2-regulated genes (D, Methods) are colored by family membership. All 78 candidates are listed in Figure S2F.

Of these 20 factors, *ZIC2* has previously been shown to be required for anterior neural tube closure in the mouse embryo ^54,55^. Before proceeding with a comprehensive knockdown screen of all 20 factors, we tested the role of *ZIC2* in regulating anterior neural tube closure in our organoid model. To knock down *ZIC2* using CRISPR interference (CRISPRi), we constructed a dual-guide RNA lentiviral transfer plasmid consisting of the top two *ZIC2-*targeting guide RNA sequences based on a published CRISPRi database ^56^ and a control plasmid consisting of two non-targeting scramble guide RNA sequences. The guide RNA sequences on each plasmid were preceded by a mouse and human U6 promoter, respectively, and followed by a distinct direct capture sequence for single-cell RNA sequencing ^57^. Each plasmid also carried a constitutive promoter driving mCerulean (Figure S2B). This dual-guide RNA construct is hereafter referred to as a ‘guide’ for brevity. In parallel, we generated a CRISPRi hPSC line in the H1/WA01 background expressing dCas9-KRAB under a constitutive promoter and confirmed that the CRISPRi construct was functional in the pluripotent state using *OCT4*-targeting guides (Figure S2C). We transduced CRISPRi hPSCs with lentivirus carrying either a *ZIC2-*targeting guide or a control non-targeting scramble guide. We differentiated lentivirus-transduced CRISPRi hPSCs into organoids as before, with two days of TGFß/WNT/BMP inhibition followed by four days of BMP4 exposure. All organoids transduced with a *ZIC2*-targeting guide (n=6) failed to undergo neural tube closure and displayed a persistent U-shaped neural plate, while all organoids transduced with a control scramble guide (n=6) formed closed neural tubes (Figure 2B). Thus, our organoid model recapitulated neural tube defects observed in *Zic2*-null mice, suggesting that *ZIC2* may similarly regulate anterior neurulation in humans.

To identify the genes regulated by *ZIC2* during neurulation, we performed single-cell RNA sequencing of Day 4 dCas9-KRAB organoids transduced with either the *ZIC2*-targeting guide or the control scramble guide. As before, we clustered cells into neural ectoderm, neural plate border, and surface ectoderm using SMD-derived marker genes from scramble control organoids (Figure S2D). In neural ectoderm cells, *ZIC2* knockdown reduced *ZIC2* expression by 60%, confirming the efficacy of the guide (Figure 2C). The proportion of neural ectoderm cells declined, and the neural plate border population expanded, suggesting that *ZIC2* loss altered cell fate proportions (Figure S2E). Notably, within the neural ectoderm, *ZIC2* knockdown led to differential downregulation of transcription factors including *PAX2, HESX1, FEZF1, OTX2, PAX8, ZNF423,* and *MAF* and upregulation of transcription factors *TCF7L1, TOX, ZNF251,* and *SIX6*, highlighting a potential gene regulatory network in the anterior neural plate (Figure 2C). We next examined regulatory regions upstream of genes strongly downregulated in Day 4 neural ectoderm cells upon *ZIC2* knockdown. Motif analysis revealed enrichment for binding sites associated with several transcription factor families, including ZNF, NR, SOX, and HES (Figure 2D, Methods). Five transcription factors from these families (*ZNF521, NR6A1, SOX2, SOX11,* and *HESX1*) appeared among the 20 transcription factors chosen for our knockdown screen based on their specific expression in the neural plate (Figure 2A, 2E, S2H), further implicating them as candidates alongside ZIC2 in co-regulating anterior neural tube closure.

We next sought to knock down our 20 candidate transcription factors in a high-throughput screen to test their roles in regulating neurulation using our organoid model. To serve as a contrasting control set for our selection criterion, we randomly selected an additional 58 transcription factors with medium to low neural specificity and a range of mean expression levels, giving us a total of 78 transcription factor candidates for our knockdown screen (Figure 2E, S2F).

### High-titer lentivirus production in small volumes enables high-throughput arrayed genetic screens in micropatterned hPSCs

Previous genetic screens in organoids have used pooled perturbations targeting multiple genes per organoid, creating mosaic tissues in which different cells carry distinct genetic modifications ^20^. Although subsequent transcriptomic analyses in such studies revealed gene-specific effects on the transcriptional states of individual cells (Figure 3A, top), this mosaic approach is ill-suited for studying morphogenesis, which is driven by coordinated changes in cellular behavior across a tissue. First, localized genetic perturbations cannot be reproducibly targeted to areas of morphological interest; second, even if the perturbation happens to localize to the region of interest, it typically is present in too few cells to influence overall tissue shape. To overcome these limitations, we developed a method to knock down single genes individually throughout entire organoids in an arrayed format, allowing us to test their roles in tissue morphogenesis (Figure 3A, bottom).

**Figure 3.**
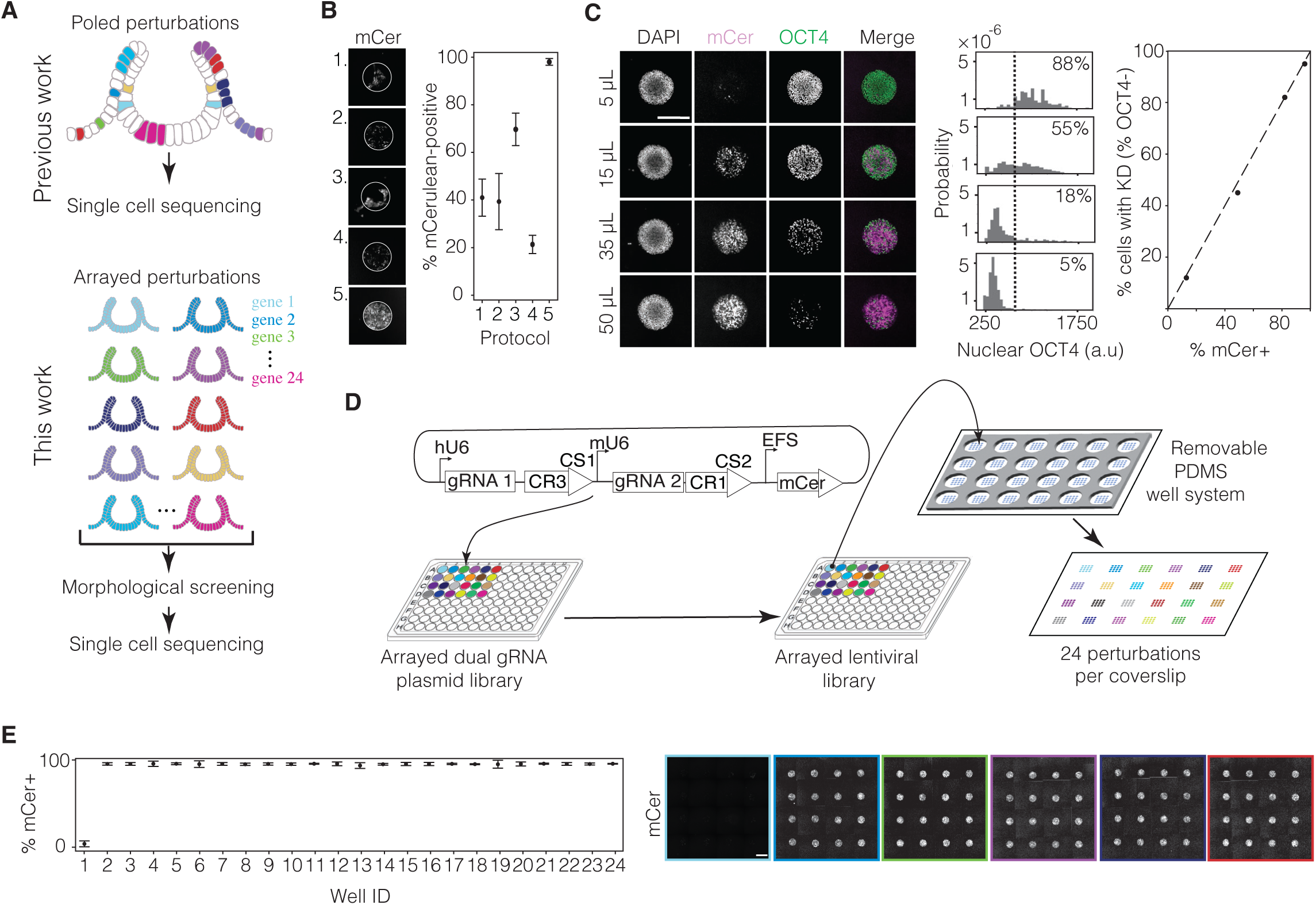
**High-throughput method for arrayed single-gene perturbations.** (A) Top: Schematic of pooled perturbations (as generated in previous work) that result in a mosaic organoid, in which a distinct sub-population of cells are perturbed by each guide RNA (shown in different colors). These mosaic organoids can be dissociated for single-cell transcriptomics. Bottom: Schematic of arrayed perturbations (generated in this work) that result in homogeneously perturbed organoids: all cells in each set of organoids are perturbed by the same guide (distinct guides shown in different colors). These organoids can be screened for morphological defects and subsequently dissociated and pooled for single cell transcriptomics. (B) Efficiency measurements of transduction protocols. Representative images (left) and quantifications (right) correspond to the following protocols. Protocol 1: Virus added 24 hours after seeding and incubated for 24 hours. Protocol 2: Virus added 24 hours after seeding and incubated for 1 hour. Protocol 3: Virus added 1 hour after seeding and incubated for 24 hours. Protocol 4: Virus added 1 hour after seeding and incubated for 1 hour. Protocol 5: Virus added during seeding and incubated for 1 hour. See Figure S3D for transduction efficiency calculation. (C) Correlation measurement of transduction and knockdown efficiencies. Left: Representative images of circular micropatterns show increasing transduction efficiency (mCerulean, mCer) and decreasing OCT4 expression with increasing viral volume of virus (volumes displayed left of each row) carrying a transfer plasmid with dual guides targeting OCT4, added to 150,000 hPSCs during seeding. Scale bar: 300 µm. Middle: Quantification of nuclear OCT4 fluorescence of individual cells in each condition. Dotted line (threshold) indicates 95^th^ percentile value of OCT4 fluorescence in mCer-cells. Right top inset in each plot is the percentage of mCer+ cells with OCT4 fluorescence above the threshold, indicating OCT4 retention. Right: Plot of percentage of cells showing OCT4 knockdown vs. percentage of cells that express mCer expressed by the guide plasmid show a positive correlation (R^2^=0.99) demonstrating that live mCer fluorescence coverage of a micropattern is an accurate indicator of percentage of cells transduced. (D) Schematic of small-volume viral growth and 24-well organoid chip seeding. Dual gRNA plasmids are first arrayed in a 96-well plate. Different colors represent unique lentiviral transfer plasmids targeting different genes (Figure S2B). HEK cells are transfected in a 96-well plate to form an arrayed viral library, then applied to a glass coverslip containing micropatterned hPSCs (depicted with blue dots) using a temporary well system (depicted in dark gray). The well system is then removed, resulting in a glass chip with 24 sets of differently perturbed organoids, each with multiple replicates, that can be cultured in a single media condition. (E) Left: Quantifications of mCer fluorescence coverage of micropatterns in a mock-transduced well without virus (Well 1, 5% median mCer fluorescence coverage, n=16 micropatterns per well) and 23 scramble vector-transduced wells (Wells 2-24, 99% median mCer fluorescence coverage across wells, n=16 micropatterns per well). See Figure S3D for calculation. Right: Stitched images of viral transduction of micropatterns in Wells 1-6. Transduction efficiency across wells is high and consistent, with a low cross-contamination rate. Scale bar: 300 µm.

To perform high-throughput arrayed knockdowns, we employed CRISPRi using a lentiviral library of dual-guide RNA constructs as above ^57^. To select guides against each target, we selected the top two targeting guides based on a published CRISPRi database ^56^. To enable an arrayed screen, we modified the cloning workflow to perform insert annealing, ligation, and transformation in parallel. Before ligation, we applied stringent gel purification following dephosphorylation of the double-digested backbone to minimize backbone contamination, which yielded a high fraction of correctly ligated plasmids (Figure S3A). As a result, we were able to directly perform minipreps from bulk-transformed bacteria cultures without colony picking, thus streamlining the cloning process. Using this protocol, we successfully generated a plasmid library of dual-guide RNA constructs targeting 77 (including *ZIC2*) out of 78 transcription factor candidates, plus a scramble control.

Achieving high-efficiency lentiviral transduction of cells requires high concentrations of lentivirus. Existing protocols to generate lentivirus, which rely on ultra-centrifugation, PEG-based concentrator, or syringe filtration ^58–60^ are laborious and not amenable to high-throughput workflows. We thus developed an approach to enable efficient, single-gene knockdowns in organoids at high throughput. We first optimized 96-well plate virus production in HEK 293T cells by testing various transfection protocols, and we found that reducing media volumes during transfection and during viral production from HEK cells significantly increased viral titers (Figure S3B-C). We then compared hPSC transduction protocols on micropatterns. Standard protocols (Sigma Aldrich) recommend adding virus 24 hours after seeding and incubating the cells with virus overnight. Using the percentage of the micropattern showing mCerulean fluorescence 48 hours post-transduction (Figure S3D) as a readout, we determined that virus treatment during cell seeding was far more efficient than virus treatment after seeding. We also found that one hour of virus treatment was sufficient to achieve high transduction efficiency. In contrast, 24 hours of high-titer virus treatment led to cell death (Figure 3B).

We hypothesized that enhanced transduction during cell seeding reflected increased exposure of the virus to the basolateral cell membrane of hPSCs. To test this, we transduced 3-day-old hPSC colonies with lentivirus after the formation of tight junctions and found that only colony edges were transduced. In contrast, transducing dissociated hPSCs during seeding led to uniform transduction (Figure S2E). These results are consistent with previous studies showing that the lentiviral VSV-G envelope protein interacts with the mammalian LDL receptor (LDLR), which is localized to the basolateral membrane of various epithelial tissues ^61^. This highlights the importance of performing viral transfection before the formation of tight junctions.

Our optimized protocol consistently achieved near-complete mCerulean expression of hPSCs on micropatterns. To determine whether mCerulean fluorescence coverage across a colony translated into efficiency of guide delivery at the level of individual cells, we cloned a transfer plasmid carrying a guide targeting the pluripotency factor OCT4 to measure knockdown in undifferentiated hPSCs. We found that 50 microliters of virus grown in a 96-well plate of HEK 293T cells, when mixed with 50 microliters of 150,000 CRISPRi hPSCs, was sufficient to deplete OCT4 in 95% of cells 48 hours post-transduction (Figure 3C). The percentage of cells showing OCT4 knockdown was highly correlated with mCerulean fluorescence coverage of the micropatterns (Figure 3C), confirming that mCerulean serves as a reliable live-cell reporter of transduction efficiency.

To perform arrayed knockdown of multiple genes in a single experiment, we needed to deliver distinct lentiviral plasmids, each carrying a different guide, to separate sets of micropatterned cells on the same coverslip. We therefore designed a PDMS stamp with 24 sets of circular micropattern features. Each set separated by the same center-to-center spacing of 9 mm matching the format of standard 96-well plates or multichannel pipette. We also created a removable PDMS well system that could partition the coverslip into 24 fluid-separated wells or be removed to create a single well (Figure 3D). In parallel, we produced lentiviruses in small volumes in 96-well plates, with virus in each well carrying a unique guide (Figure 3D). To seed hPSCs, we coated the entire PDMS stamp with Matrigel and transferred it onto a 50-by-75 mm glass coverslip. Following Matrigel transfer, we removed the stamp and replaced it with the PDMS well system. Using a multichannel pipette, we added a suspension of single hPSCs and lentivirus into each well (Figure 3D, Methods). After 70 minutes of incubation, we thoroughly rinsed and removed unadhered cells and unfused virus. 48 hours later, we assessed mCerulean fluorescence coverage of micropatterns. Our protocol resulted in highly reproducible transduction of micropatterned hPSC colonies in an arrayed format, with a median transduction efficiency of 99% (Figure 3E). The PDMS well system prevented viral cross-contamination between neighboring wells, as seen by the less than 5% transduction of the untreated control well. Transduced hPSC colonies could then be differentiated into anterior neural tube organoids to model the effects of single-gene perturbations on neural tube morphogenesis.

### A high-throughput genetic screen of anterior neurulation reveals three transcription factors necessary for proper neural tube closure

We cloned guide plasmids (Methods) against 77 out of our 78 transcription factor candidates (Fig. Figure 2E, S2F). Using our screening method, we conducted a high-throughput arrayed CRISPRi knockdown screen of these 77 factors to evaluate their individual roles in anterior neurulation (Figure S2E). We included *ZIC2* and a scramble control in this screen for methodological consistency with all other factors. Guide plasmids in our screen yielded a mean transduction efficiency of 93% (Figure S4A-B), slightly lower than the 99% we achieved with a clonal guide (Figure 3E), perhaps due to mis-ligated products in the non-clonal plasmid population (Methods). After seeding and transducing hPSCs, we differentiated them into organoids using our protocol (Figure S1D). Organoids were fixed after four days of BMP4 exposure, stained for NCAD and ECAD, and screened for morphological defects using epifluorescence microscopy. Analysis of guide expression at the end of the differentiation time course, as measured by mCerulean, showed retention of lentiviral gene expression throughout the experiment (Figure S4F).

We focused particularly on neural tube closure defects, as assessed by a lack of continuous NCAD staining enclosing the neural lumen. Lumen morphology was clear from epifluorescence microscopy for all candidates except *RFX7* and *ZNF521*, which required confocal microscopy for phenotypic categorization (Figure 4B, S4C). Most candidates had no closure defects (score = 0, Figure 4A-C, Figure S4D). Knockdown of 24 transcription factors (*HESX1, TGIF1, ZFHX4, MAFB, TSHXZ2, ZNF219, PAX3, ID3, PBX3, SP5, POU3F2, SSRP1, TLX2, VSX2, DNMT1, CSNRP3, TFDP2, DNMT3B, ZNF280, MSX1, SALL3, PLAGL1, SALL4, ARX*) showed minor closure defects upon knockdown, where a fraction of organoids showed either a small opening in the neural plate or delayed neural tube closure compared to control organoids (score=1, Figure 4A-C, Figure S4E). For example, organoids with a *HESX1* knockdown showed a minor opening in the neural plate on Day 4 (Figure 4C), which resolved after an additional day of differentiation in five out of six organoids (Figure 4B).

**Figure 4.**
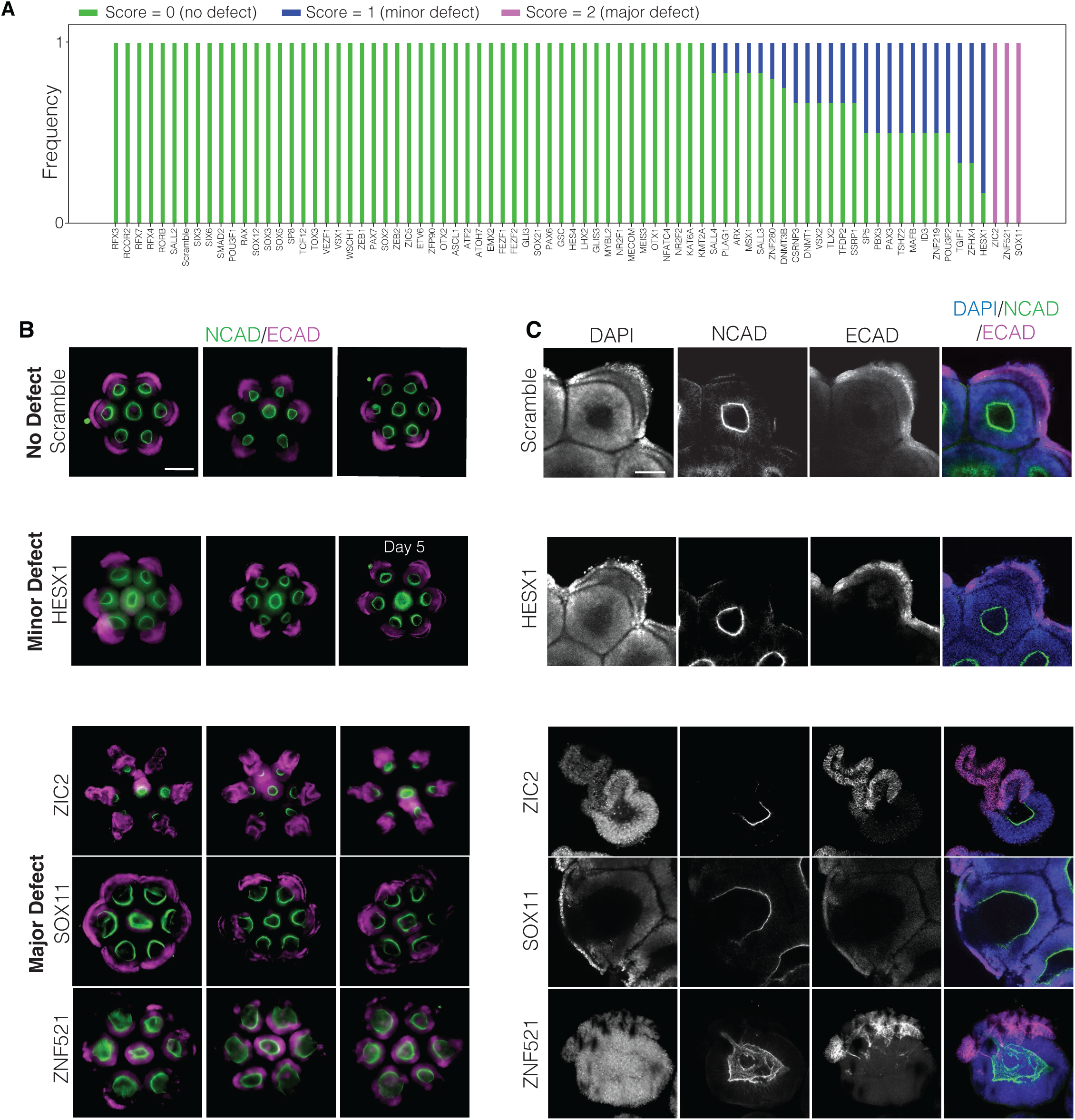
***SOX11, ZIC2* and *ZNF521* are necessary for neural tube closure** (A) Phenotypic scores of organoids knocked down with each gene. Scoring was based on whether organoids had no closure defects (score = 0, white), a minor defect (score = 1, gray), or a major defect (score = 2, black) consisting of either a fully opened neural plate or multiple closure points. (B) Epifluorescence microscopy of organoids at Day 4 (unless otherwise noted), showing examples of no defect (top row, scrambled control), minor defect scores (*HESX1* knockdown, row 2), or major defect scores (*ZIC2*, *SOX11* and *ZNF521* knockdown, rows 3-5 respectively). Three biological replicates per condition are shown with n=6 outer organoids per replicate. Imaging of *HESX1* knockdown organoids with a minor defect on Day 5 show that most of them eventually close and that minor openings represent delayed closure. Scale bar: 300 µm. (C) Corresponding confocal images of single organoids for each of the phenotype classes in (B), showing a range of closure phenotypes. Scale bar: 100 µm.

Knockdown of three transcription factors (*ZIC2*, *SOX11*, and *ZNF521*) showed major closure defects upon knockdown (score=2) consistently across all organoids in three biological replicates consisting of 6 organoids each (Figure 4A-C). Confocal microscopy of *ZIC2* and *SOX11* knockdown organoids showed that their neural plates were fully open, while confocal microscopy of *ZNF521* knockdown organoids showed multiple discontinuous regions of surface ectoderm on top of the neural ectoderm, which we interpreted as multiple points of closure (Figure 4C).

To quantify the knockdown efficacy of CRISPRi guides in organoids, we performed scRNA-seq on Day 4 organoids with arrayed knockdowns of the transcription factors that showed a major phenotype (*ZIC2, SOX11* and *ZNF521*). To determine false negative rates or minor phenotypes as a result of poor knockdown efficacy, we also sequenced organoids corresponding to knockdowns of six transcription factors that did not show a closure defect (*OTX2, PAX6, POU3F1, LHX2, TOX3*, and *GLI3*) and two transcription factors that showed a minor defect (*TGIF1* and *ZFHX4*). Because each guide included a constant region tagged with a direct capture sequence readable by scRNA-seq ^57^, we could pool the arrayed samples prior to single-cell library preparation to reduce costs (Figure S2B). Guide targets *ZIC2*, *SOX11*, and *ZNF521* were among the most downregulated transcripts within the neural ectoderm in each knockdown condition, respectively (boxed data, Figure 5A, Figure S5A-B), showing that targeted knockdown was successful. Knockdown efficacies of guides that did not show a phenotype or showed only a minor phenotype were comparable to knockdown efficacies of *ZIC2* and *ZNF521*, with 7 out of 8 chosen guides showing significant knockdown (Figure S5C).

**Figure 5.**
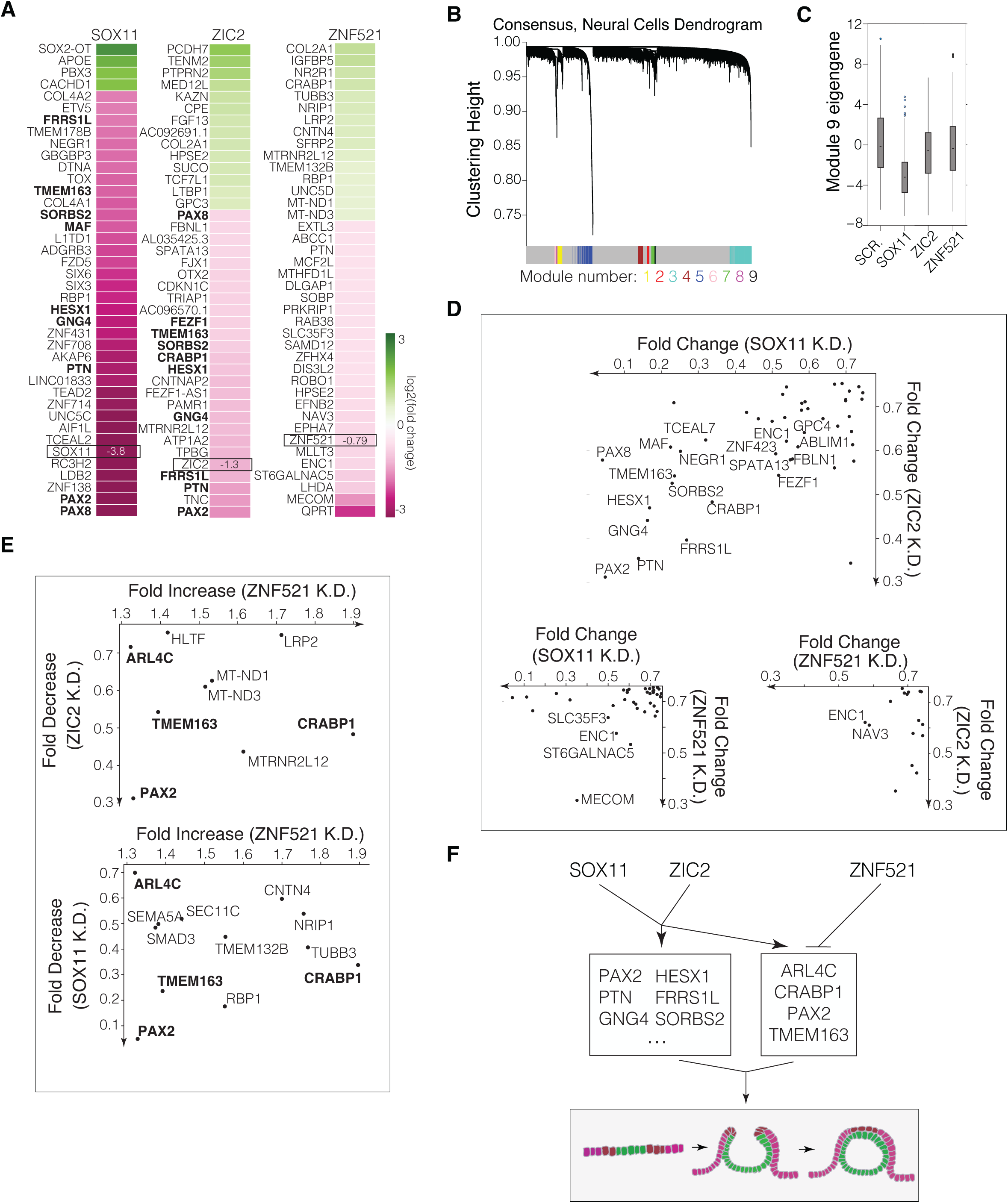
***SOX11* and *ZIC2* co-regulate a shared set of genes in opposition to *ZNF521*.** (A) Fold changes of the 40 most up- or down-regulated genes in the neural ectoderm of each knockdown condition at Day 4, as compared to organoids transduced with a scramble control guide. Genes that are co-downregulated by *SOX11* and *ZIC2* knockdown are shown in bold. (B) Cluster dendrogram of WGCNA analysis of scRNA-seq gene expression from neural cells (scramble control) showing modules of co-expressed genes. Module assignment (numbered) is indicated by the color bar; genes labeled in gray were not assigned. (C) Module eigengene values (representative first principal component of expression of genes in a module) of Module 9 per cell in the neural cell population in each knockdown condition compared to scramble control data. *SOX11* knockdown leads to significant downregulation of genes in Module 9 (p < 0.0001), which includes *ZIC2* (Fig. S2D). (D) Scatterplots showing fold changes of significantly co-downregulated genes (Anderson Darling test, p<0.05) in the neural cell populations from transcription factor knockdown pairs, as compared to organoids transduced with a scramble control guide. Genes are plotted if their fold change expression (FC) relative to scrambled control passes the threshold log2(FC) <-0.4 in both knockdown conditions. Genes are labeled if log2(FC) <-0.6 in both conditions. (E) Scatterplots showing fold changes of genes that are significantly downregulated (Anderson Darling test, p<0.05) by *ZIC2* or *SOX11* knockdown and significantly upregulated by *ZNF521* knockdown, as compared to organoids transduced with a scramble control guide. (F) Gene regulatory model: *SOX11* and *ZIC2* are upstream a similar set of genes, including neural tube defect candidates from the literature like *PAX2, HESX1*, and *CRABP1*. *ZNF521* upregulates a separate set of genes but downregulates several shared *ZIC2*- and *SOX11*-upregulated genes.

Consistent with our hypothesis that the neural plate drives neurulation, the three transcription factors whose knockdown most significantly disrupted neural tube closure (*ZIC2*, *SOX11*, and *ZNF521*) were all among the list of 20 transcription factors chosen based on specific expression in the neural ectoderm (Figure 2A). Notably, strong phenotype-producing factors *SOX11* and *ZNF521* and minor phenotype-producing factor *HESX1* all belonged to transcription factor families with associated motifs at regulatory regions near *ZIC2* target genes (Figure 2D, E). This suggests the possibility of a coordinated transcriptional program within the neural ectoderm downstream of these factors.

### Deciphering the gene networks regulating anterior neurulation

Morphologically, *ZIC2* and *SOX11* knockdowns resulted in a persistently open neural plate, whereas *ZNF521* knockdown gave rise to multiple discrete closure points, suggesting opposing roles (Figure 4B-C). Phenotypically, these results suggest that *ZIC2* and *SOX11* promote neural tube closure, while *ZNF521* has an opposing role. To determine whether the phenotypic similarities and differences of *ZIC2*, *SOX11*, and *ZNF521* knockdowns are reflected in the gene regulatory functions of these factors, we analyzed single-cell gene expression data from perturbed organoids.

We analyzed scRNA-seq data from Day 4 organoids in which each of these three transcription factors were knocked down to evaluate changes in gene expression relative to control organoids. Hierarchical clustering of genes expressed in neural ectoderm from control organoids revealed nine modules of co-regulated genes (Figure 5B). One module, consisting of *ZIC2* and several other neural transcription factors and markers, was significantly downregulated following *SOX11* knockdown (Figure 5C, S5D), consistent with the enrichment of *SOX* family binding motifs at regulatory regions near *ZIC2*-regulated genes (Figure 2D).

We next examined genes that were significantly downregulated in the neural ectoderm in at least two of three knockdowns of *ZIC2, SOX11* and *ZNF521,* compared to the scramble control (Anderson-Darling test, p < 0.05, Figure 5D). Of these genes, those affected by both *SOX11* and *ZIC2* knockdown showed larger fold decreases than all other pairs of knockdowns, as reflected by the relative number of genes in the lower left quadrants of fold change plots comparing knockdown pairs (Fig. 5D, scatterplots). Many of these genes also ranked highly in their degree of downregulation when compared to all other genes upon *SOX11* or *ZIC2* knockdown (Fig. 5A).

These observations suggested that SOX11 and ZIC2 jointly drive a gene regulatory program consisting of a shared set of genes, including *PAX2, PTN, GNG4, HESX1, FRSS1L, CRABP1, SORBS2, TMEM163, PAX8,* and *NEGR1* (from bottom left to top right in Figure 5D). Of these genes, *PAX2* knockouts show fully penetrant exencephaly, although at the midbrain level rather than forebrain ^62,63^, and *CRABP1* single-nucleotide polymorphisms have been found in human patients with spina bifida and exencephaly ^64^.

Consistent with the opposing role of *ZNF521* suggested by its knockdown phenotype, several genes that were significantly (p < 0.05) downregulated upon *ZIC2* or *SOX11* knockdowns were upregulated upon *ZNF521* knockdown (Fig 5E). Four of these genes, including neural tube literature candidates *PAX2* and *CRABP1*, were significantly downregulated upon both *ZIC2* and *SOX11* knockdown and significantly upregulated upon *ZNF521* knockdown (*PAX2, CRABP1*, *TMEM163,* and *ARL4C*; Figure 5E).

We next asked whether we could phenocopy a neural tube closure defect by knocking down genes downstream of *ZIC2*, *SOX11* and *ZNF521*. We selected ten genes that were most co-downregulated by *SOX11* and *ZIC2,* including *PAX2, PTN, GNG4, HESX1, FRSS1L, CRABP1, SORBS2, TMEM163, PAX8,* and *NEGR1* (Fig. 5D). We also chose *CRABP1, PAX2, TMEM163,* and *ARL4C,* which were significantly downregulated upon *SOX11* and *ZIC2* knockdown and significantly upregulated upon *ZNF521* knockdown (Figure 5E). Apart from *HESX1,* which showed a minor closure defect (Figure 4A-C), there was no single gene in this set whose knockdown alone recapitulated the open neural plate phenotype observed upon *ZIC2* or *SOX11* knockdown (Figure S5F). These findings argue that *ZIC2*, *SOX11*, and *ZNF521* do not act through a single dominant downstream effector and instead regulate a combination of genes to drive anterior neurulation.

## Discussion

Perturbing the expression of genes individually and uniformly in hPSC-derived tissues is essential to determine their roles in morphogenesis during human embryonic development. The arrayed CRISPRi platform developed in this study allowed us to screen transcription factors in an hPSC-derived organoid model of anterior neurulation, revealing that three key transcription factors, *ZIC2, SOX11*, and *ZNF521,* act together to drive neurulation in the forebrain region. Knocking down these transcription factors shows opposing phenotypes, with *ZIC2* and *SOX11* knockdown resulting in an open neural plate and *ZNF521* knockdown leading to multiple points of neural tube closure. Our analyses of gene expression changes upon *ZIC2* and *SOX11* knockdown suggest that these two transcription factors regulate a shared set of gene targets. Consistent with the opposite phenotype of *ZNF521*, we find that *ZNF521* downregulates several of the gene targets upregulated by both *ZIC2* and *SOX11*. The set of these oppositely regulated factors include *PAX2* and *CRABP1,* which have been previously implicated in anterior neural tube defects in mice and humans, respectively ^62–64^.

Our results suggest that coordinated transcriptional regulation of downstream gene targets by *ZIC2*, *SOX11*, and *ZNF521* are necessary for successful anterior neurulation *in vitro* (Fig. 5F). The role of *ZIC2* in neurulation has been well characterized *in vivo* in mouse ^54,55^ and genetic variants of *SOX11* in humans have been associated with midline defects, microcephaly and ocular malformations ^65,66^. In mouse, *SOX11* deficiency leads to grossly normal neurulation in mice despite causing other broad morphological defects, suggesting possible species-specific differences in *SOX11*’s developmental roles ^67^. As human anencephaly is embryonic lethal, genetic causes have only been obtained in rare cases with whole exome sequencing ^68,69^. This raises the possibility that *ZIC2*, *SOX11*, and *ZNF521* also drive neurulation *in vivo*. In the future, it will be of interest to identify more genes with *de novo* mutations contributing to anencephaly in human embryos, as has recently been conducted for spina bifida ^70^.

The ability to perform single-gene perturbations in robust *in vitro* models of human development is essential for dissecting the mechanisms that drive human embryonic morphogenesis. Until now, this goal has been limited by variability in organoid morphogenesis and the lack of scalable methods for uniform single-gene perturbations. The platform we report here integrates a micropatterned hPSC-based protocol to generate reproducible organoids with uniform single-gene knockdown via lentivirus-mediated arrayed CRISPR interference to identify genes with roles in morphogenesis. Furthermore, the method reduces costs compared to clonal knockdown approaches, is scalable, and can be performed in an academic laboratory setting, enabling tissue-wide single-gene perturbations at a scale that has not been previously feasible either in organoids *in vitro* or in mammalian models *in vivo*. Our approach bridges a critical gap between the genetic study of traditional model organisms and human developmental biology, offering a path for new mechanistic insights and the discovery of therapeutic targets for neural tube defects and other congenital malformations.

## Limitations of the study

The first limitation of this study is that inherent to any method that utilizes CRISPR: the efficacy of selected guides. Based on quantification of gene knockdown by scRNA-seq, we estimate that 87% of targeting guides were effective in our system. The second limitation is the assumption that the genes we discovered *in vitro* are relevant in humans *in vivo*. False positives or false negatives could arise in part due to the study of isolated tissues *in vitro* that lack surrounding cell types, which may either compensate for or trigger tissue-level defects. Given the increasing fidelity of organoid models in terms of their ability to recapitulate the complexity of fetal tissues, we believe that this limitation will be overcome in time. While *in vitro* systems are ultimately models, they serve as a complement to *in vivo* studies to increase our understanding of human development and disease, and they offer value in their amenability to high-throughput perturbations.

## Acknowledgements

We thank Nicole Ramirez, Claire Reardon, Jeffery Nelson, Claire Maesner, Christian Daly, Nathan Weeks, and Zachary Niziolek at the Bauer Core Facility at Harvard University for performing single-cell RNA sequencing and assisting with FACS for cell line generation, and Doug Richardson at the Harvard Center for Biological Imaging for help with imaging. We thank Cassandra Extavour, Sean Megason, Jessica Whited, Vlad Denic, and members of the Ramanathan lab for scientific discussions and comments. This work was supported by NIH R01MH136014 (SR), NIH R01GM131105 (SR), and startup funds from Harvard University. R.E.H. was additionally funded by a National Science Foundation Graduate Research Fellowship.

## Author Contributions

R.E.H., G.M.A., and S.R. conceptualized and designed the study. R.E.H., G.M.A., and H.C.M. performed the experiments. R.E.H., G.M.A., and J.C. performed the single cell analysis. R.E.H. and C.A. performed image analysis. R.E.H., G.M.A., and S.R. wrote the manuscript.

## Declaration of Interests

S.R., G.A., and R.H. are authors on the following patent application, which contains aspects of this work: “Bioengineering and machine learning framework for complex tissue development.” Application serial number PCT/US24/28838.

---

**Figure S1.**
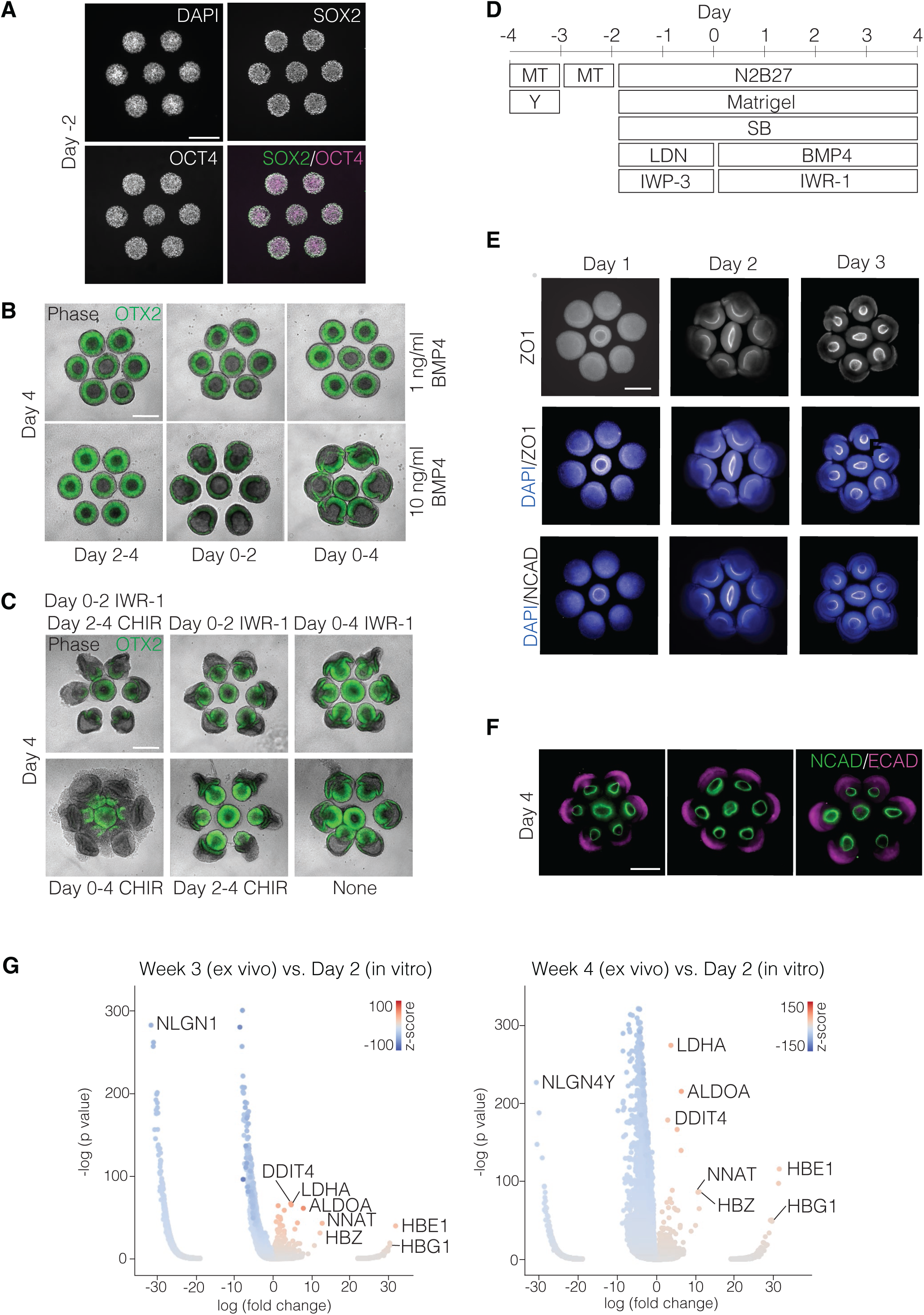
**Development and validation of a hPSC-derived *in vitro* model of anterior neurulation.** A) Epifluorescence images of fixed and stained hPSCs 2 days after seeding (Day −2 according to scheme in S1D) onto hexagonally arranged 250-µm diameter micropatterns with 100-µm spacing (Methods), showing co-expression of pluripotency markers SOX2 and OCT4. Scale bar: 300 µm. B) Live epifluorescence images of neural tube organoids expressing an OTX2-Citrine reporter to evaluate responses to different BMP4 treatment courses. The bottom right condition (Day 0-4 BMP4, 10 ng/ml) was selected for further experiments. Scale bar: 300 µm. C) Live epifluorescence images of neural tube organoids expressing an OTX2-Citrine reporter to evaluate responses to different WNT activation and inhibition courses with agonist CHIR99021 (CHIR) and inhibitor IWR-1-endo (IWR-1). The top right condition (Day 0-4 IWR-1, 1 µM) was selected for further experiments. Scale bar: 300 µm. D) Neural tube organoid differentiation protocol. hPSCs are seeded onto 2D micropatterns on Day −4 (2 days prior to differentiation) in mTeSR Plus (MT) with 10 µM ROCK inhibitor Y-27632 (Y). Media is changed to MT without Y on Day −3. On Day −2, organoids are folded into 3D cysts in high-concentration Matrigel and simultaneously differentiated toward ectodermal fates in N2B27 media (Methods) by addition of 10 µM TGFB inhibitor SB431542 (SB), 0.5 µM BMP inhibitor LDN193189 (LDN), and 1 µM WNT inhibitor IWP-3 on Day −2. On Day 0, addition of 10 ng/ml BMP4 induces mediolateral patterning of the ectoderm, while WNT inhibition is sustained by 1 µM IWR-1 addition. BMP4-containing media is replaced on Day 2. Each media replacement includes fresh high-concentration Matrigel. E) Epifluorescence images of organoids after 1, 2, and 3 days of BMP4 treatment, fixed and stained for neural ectoderm marker NCAD, tight junction marker ZO1, and nuclear marker DAPI. Scale bar: 300 µm. F) Epifluorescence images of three biological replicates of neural tube organoids on Day 4, showing complete closure in the form of a continuous ring of NCAD. 100% of organoids show closure (see Figure 1D). Scale bar: 300 µm. G) Volcano plots showing neural genes that are differentially expressed between Week 3 embryos and Day 2 organoids (left) and Week 4 embryos and Day 2 organoids (right). The x-axis refers to the log(fold change) of mean single-cell gene expression in embryos divided by mean single-cell gene expression in organoids. The color bar shows the score by Scanpy’s t-test across two groups.

**Figure S2.**
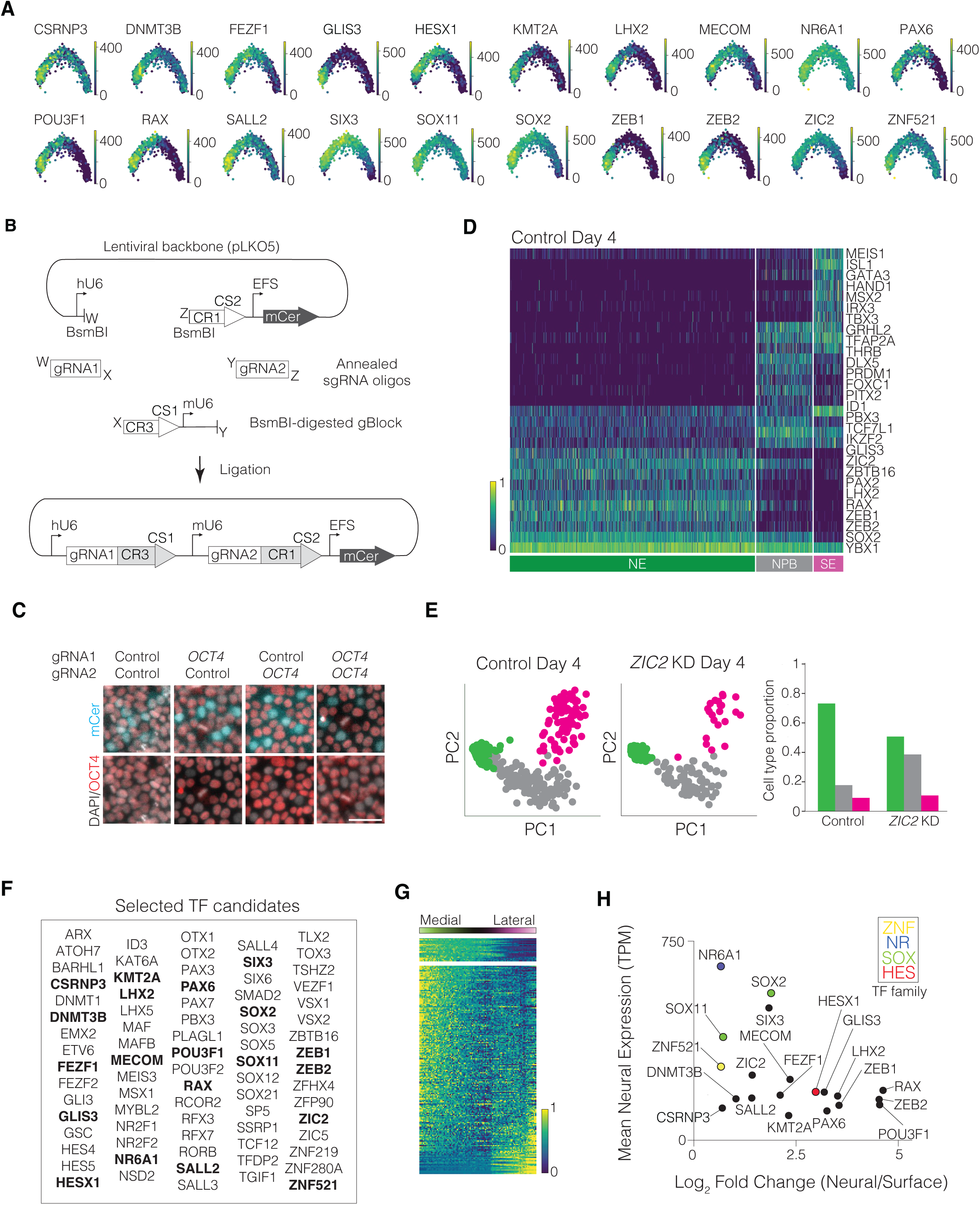
**Transcriptome-based identification of candidate transcriptional regulators of anterior neurulation.** A) Principal component analysis of single-cell RNA-sequencing data from Day 2 organoids, colored by expression of 20 most medially expressed transcription factors. Gene expression is transcript count-normalized, and units are in transcripts per million. Scale bar for each sub-panel is shown to the right of the PCA plot. B) Cloning schematic for lentiviral dual-guide RNA vector (based on Replogle, *et al*., 2020), for which pLKO5 backbone was modified to carry an mCerulean (mCer, a cyan fluorescent protein) reporter, digested with BsmBI, and ligated with two pairs of annealed sgRNA oligos together with a BsmBI-digested gBlock. The final vector encodes gRNA1-CR3-CS1 under a human U6 (hU6) promoter, gRNA2-CR1-CS2 under a mouse U6 (mU6) promoter, and mCerulean under an EFS promoter (Methods). C) dCas9-KRAB CRISPRi H1 (WA01) transduced with dual-guide RNAs consisting of either a control non-targeting scrambled sequence (“control”) or *OCT4*-targeting sequence (“*OCT4*”) under the hU6 or mU6 promoter in guide RNA positions 1 (“gRNA1”) and 2 (“gRNA2”), respectively, fixed and stained for DAPI and OCT4 48 hours post-transduction. Columns correspond to a control sequence at both positions (first column), an OCT4 sequence in position 1 and control sequence in position 2 (second column), a control sequence in position 1 and OCT4 sequence in position 2 (third column), and an OCT4 sequence at both positions (fourth column). Top and bottom rows correspond to merged channel images of the same fields of cells stained for OCT4 (red) and DAPI (gray), with or without the mCerulean channel (cyan) overlaid. The cells transduced with virus (mCer+) in the second, third, and fourth columns, but not those the first column, have low OCT4 levels, demonstrating that the *OCT4*-targeting guide is specific and functional when present in the first, second, or both positions. Scale bar: 50 µm. D) scRNA-seq of Day 4 organoids transduced with a control non-targeting scramble dual-guide RNA construct. Control cells were clustered as neural ectoderm (NE, green), neural plate border (NPB, gray), or surface ectoderm (SE, magenta) based on expression of labeled transcription factors. Gene expression (color bar, bottom left) is normalized to median transcript count, followed by log- and min-max normalization. E) Left: Cells from scRNA-seq of Day 4 scramble control and *ZIC2* knockdown organoids plotted along the first two principal components for control cells. The data points corresponding to the three cell types are colored appropriately (NE, green; NPB, gray; SE, magenta). Right: Histogram of cell type proportions shows that *ZIC2* knockdown organoids have less NE, more NPB, and a similar SE fraction compared to control organoids. F) Full list of 78 transcription factor candidates selected for knockdown, with 20 neural transcription factors selected based on mediolateral expression (Figure 2A, S2A) in bold. G) Standardized mediolateral expression profiles of top 200 most highly expressed transcription factors in Day 2 organoids, ordered from most medially expressed (top) to most laterally expressed (bottom). The 20 most medially expressed transcription factors (Figure 2A, right) are separated above the rest. H) Plot of mean expression in neural ectoderm (transcripts per million, TPM) versus log fold change in expression between the neural and surface ectoderm in Day 2 organoids for the top 20 mediolaterally enriched transcription factors from Figure 2E. For all 20 transcription factors, mean expression > 70 TPM and log2(fold change) > 0.6.

**Figure S3.**
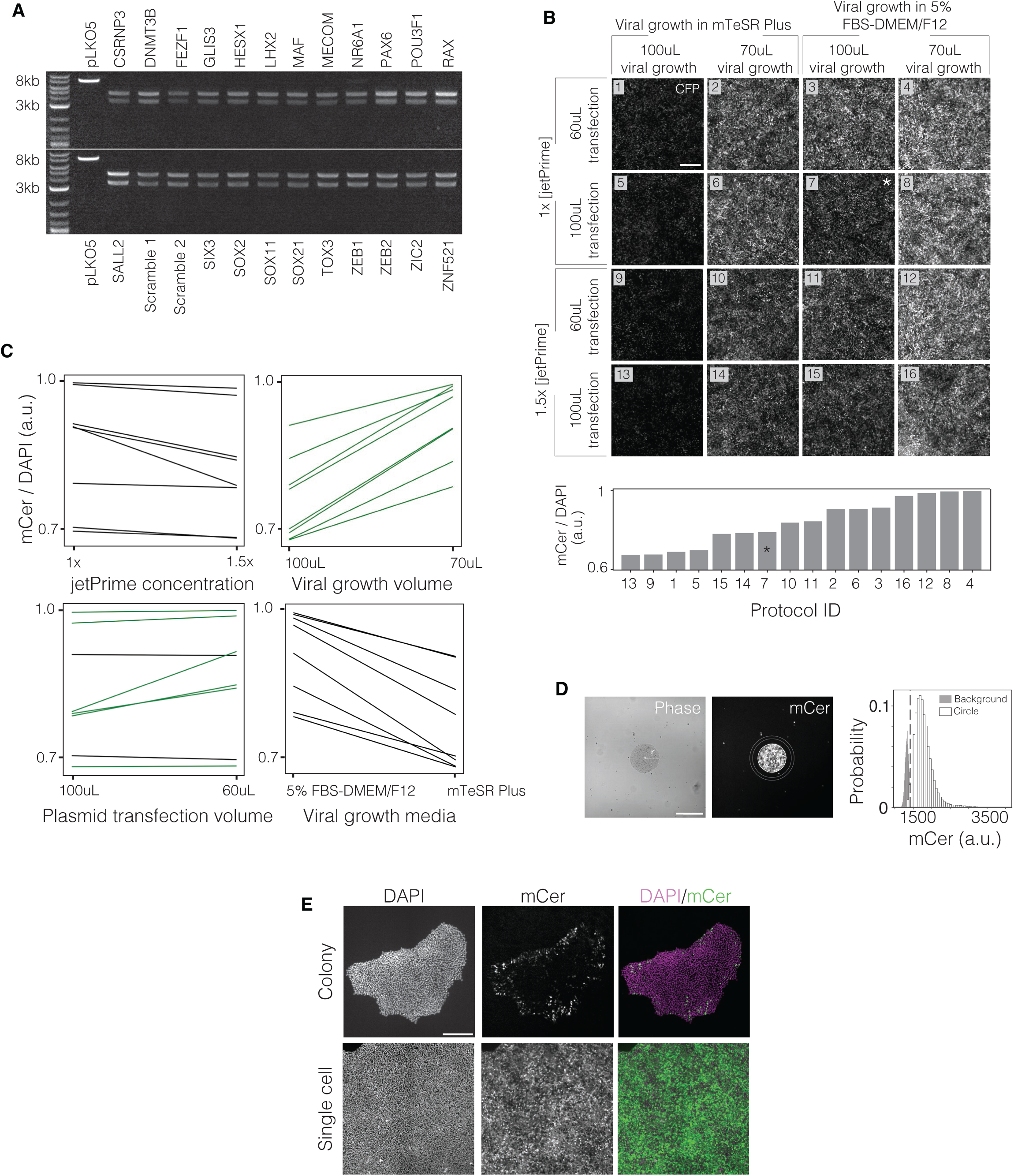
**Development of transfection and transduction methods.** (A) Gel electrophoresis of non-clonal plasmid stocks (see Figure S2B, Methods) after double digest with BamHI and DraIII. Expected band length for backbone (control lane): 7.4kb. Expected band lengths for backbone with ligated inserts (following lanes): 3.4kb and 4.5kb. Ladder: NEB 1kB Plus. Band intensities show that a majority of non-clonal plasmid stock contains the desired insert. (B) Top: Epifluoresence images of mCerulean (mCer) expression in confluent hESC culture, 48 hours after lentiviral transduction with scramble control vector from one of 16 different HEK transfection protocols. Protocols altered 4 variables in a 96-well transfection of HEK cells. Bottom: Ratio of mCer to DAPI signal from each protocol above, rank ordered by transduction efficiency. Protocol 7 (*) is the standard protocol recommended by jetPrime. Scale bar: 300 µm. (C) Ratio of mCer to DAPI signal from Figure S3B, paired by shared variables to show the effect of changing the variable in each controlled pair. Changes that resulted in increased transduction are shown in green. General trends show that decreasing viral growth volume and decreasing plasmid transfection volume result in higher viral production. (D) Example of transduction efficiency analysis. Left: Colony diameter as measured by colony segmentation by Ilastik in the phase channel. Middle: Colony segmentation as applied to the mCer channel (white) and 2D background halo (area between gray circles). Right: Quantification of mCer signal from colony (white) and background halo (gray). Dotted line (threshold) represents 95^th^ percentiles of mCer expression in the background halo. This approach is validated with nuclear analysis in Figure 3C. Scale bar: 300 µm. (E) mCer expression in hESC colonies, 48 hours after transduction with scramble control vector made with Protocol 4 from Figure S3B. Top: hESCs were transduced after 48 hours of adherent growth on plates. Bottom: hESCs were transduced in single cell suspension directly after dissociation. Scale bar: 300 µm.

**Figure S4.**
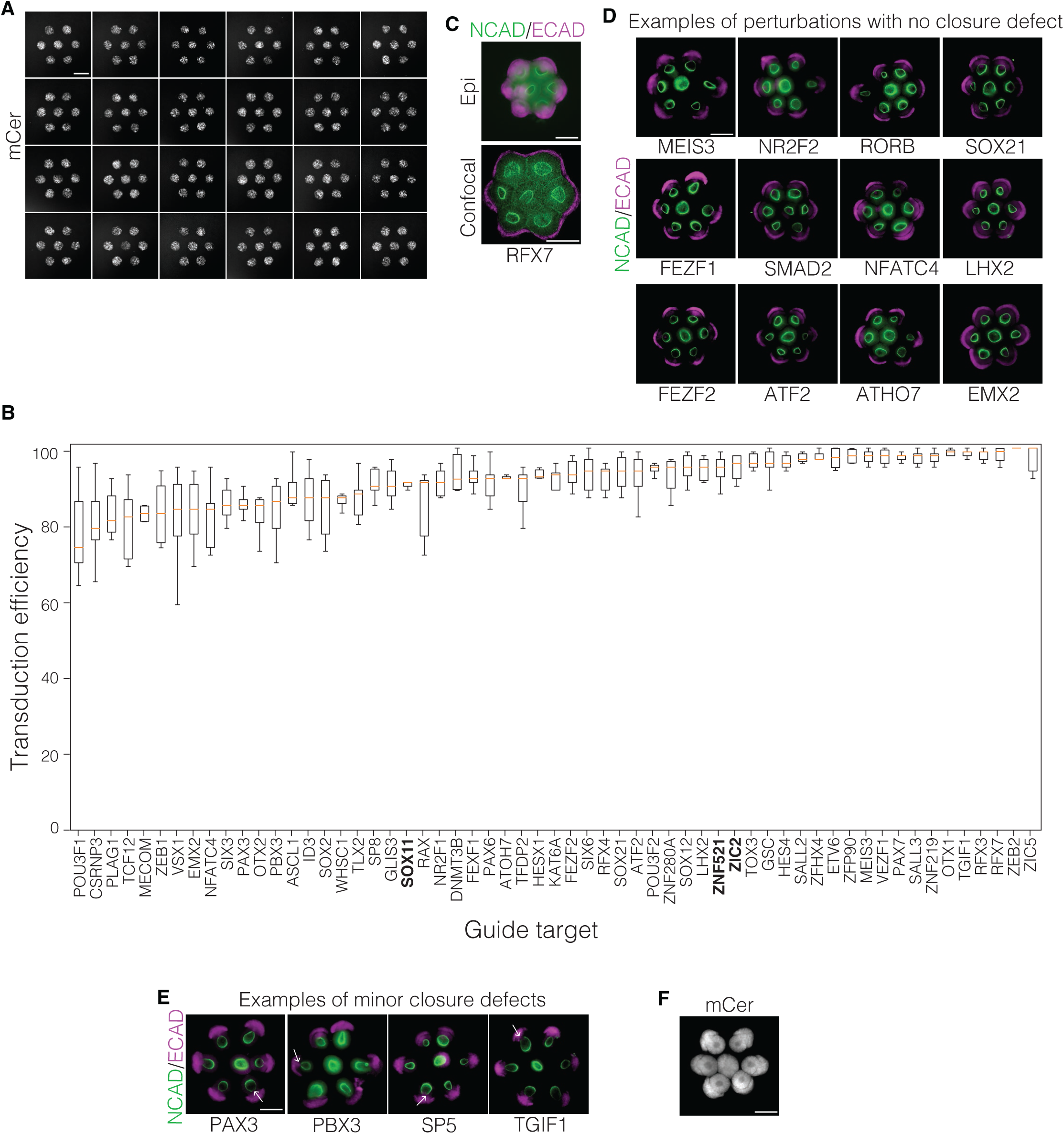
**Perturbation of most transcription factors does not give rise to morphological phenotypes.** (A) 24 examples of live epifluorescence microscopy of lentiviral marker mCerulean (mCer), 48 hours after transduction in suspension and seeding on micropatterns as in Figure 3D. Scale bar: 300 µm. (B) Quantification of transduction efficiencies from individual sgRNAs, 48 hours after transduction in suspension and seeding on micropatterns as in Figure 3D. The median transduction efficiency across all guides using the non-purified plasmid prep was 93%. (C) Top: Epifluorescence image of *RFX7* perturbation. Bottom: Maximum intensity projection of a confocal Z-stack (bottom). All organoids received a score of 0 (no phenotype) after confocal imaging. Scale bar: 300 µm. (D, E) Epifluorescence microscopy of terminal organoid morphologies, differentiated as in Figure S1D for 6 days, then fixed and immunostained. E: Examples of minor openings are indicated in white arrows. Scale bar: 300 µm. (F) Distribution of mCer in organoids transduced with a scramble guide, measured 8 days post-transduction (2 days of mTeSR growth and 6 days of differentiation). Scale bar: 300 µm.

**Figure S5.**
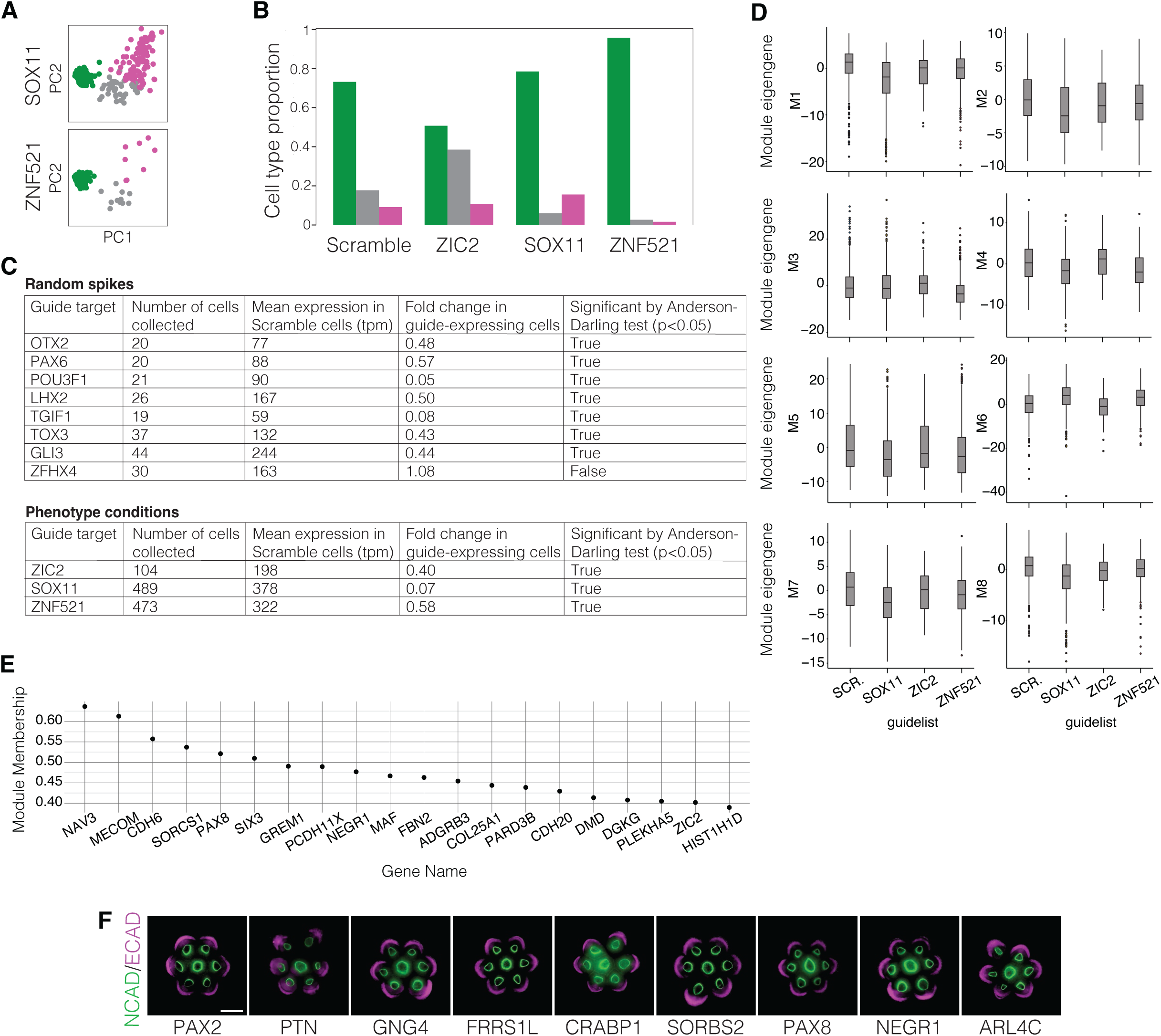
**Analysis of scRNA-seq data from perturbed organoids.** (A) Principal component analysis of cell types in Day 4 organoids infected guides against *SOX11* or *ZNF521*. Cells are plotted using the two major principal components from the scramble control data from Figure S2E. Cell type assignments to neural ectoderm (green), neural plate border (gray), and surface ectoderm (magenta) are based on expression of canonical transcription factors from scramble control data in Figure S2D. (B) Quantification of cell type proportions from scRNA-seq of all control and major closure defect datasets (*ZIC2*, *SOX11*, and *ZNF521* knockdown). Cell types are assigned as in (A). (C) Top: Knockdown efficacy calculations of sgRNAs that led to no phenotype or minor closure defects. Seven out of eight guides showed significant knockdown of their target gene with a knockdown efficacy of >40%. Bottom: Knockdown efficacy calculation of sgRNAs that produced a major closure defect. All three guides showed significant knockdown of their target gene with a knockdown efficacy of >40%. (D) Dendrogram module eigengene values from the remaining modules from Figure 5C. No other module shows significant changes in expression in any perturbation condition tested. (E) Correlation values with module eigengene (module membership) of top 20 genes in Module 9, including ZIC2, from Figure 5C. (F) Epifluorescence images of Day 4 organoids perturbed with guides against downstream targets of *SOX11*, *ZIC2*, and/or *ZNF521*. Scale bar: 300 µm.

**Supplementary Video 1.** Timelapse of actin during neural folding, hours 0-7 after BMP4 addition. Timestep = 30 minutes.

**Supplementary Video 2.** Timelapse of actin during neural folding, hours 9-33 after BMP4 addition. Timestep = 60 minutes.

## DATA AND RESOURCE AVAILABILITY

Raw scRNA-seq data will remain unpublished to protect donor patient privacy, in accordance with Institutional Review Board (IRB) protocol and updated WiCell contract restrictions on publishing RNA sequences derived from human embryonic stem cells. Cell-by-gene count matrices from this paper are available at Figshare dataset 2026_HuangAnand. Code from this paper is available at GitHub repository royahuang/2026_HuangAnand. Any requests for plasmids and cell lines will be fulfilled by the lead contact, Sharad Ramanathan. Materials will be provided upon completion of a Material Transfer Agreement.

## METHODS

### Cell lines

Neural tube organoid differentiation was optimized in an H1 OTX-YFP hESC line (made at and karyotyped by the Allen Institute). Knockdown experiments were conducted with an H1 DCX-YFP hESC line (made at and karyotyped by the Allen Institute) modified by inserting pAAVS1-NDi-CRISPRi into the AAVS1 locus. pAAVS1-NDi-CRISPRi (Gen1) was a gift from Bruce Conklin (Addgene plasmid #73497; http://n2t.net/addgene:73497; RRID:Addgene_73497). dCas9 integration was validated with genotyping and CRISPRi activity was validated with OCT4 knockdown. The H1 background line has XY chromosomes and was chosen because of our extensive experience in the lab with this line. We recognize the value of studying lines with other genotypes in order to learn sex-specific mechanisms of human development. As there is considerable line-to-line variability in stem cell models^71^, this work provides one benchmark against which to compare XX other XY lines. All research in this study falls under International Society for Stem Cell Research Category 1a.

hESCs were grown in 6-well culture dishes (CellTreat) pre-treated with diluted Matrigel hESC-qualified Matrix (Corning Cat No. 354277) according to manufacturer’s instructions and fed with mTeSR Plus (StemCell).

hESCs were maintained via colony-passage every four days with ReLeSR (StemCell) according to manufacturer’s instructions. Colonies were broken into clumps of 5-15 cells and diluted 1:30 into a new maintenance plate using wide-bore pipette tips. Cells were fed with fresh mTeSR Plus every two days. Cell lines were routinely tested for mycoplasma contamination (Mycoplasma PCR Detection Kit, ABM G238).

hESCs were expanded for experiments via single cell passage to ensure consistent cell density. 10cm culture dishes were pre-treated with Matrigel according to manufacturer’s instructions. Maintenance hESC wells were rinsed with PBS, treated with Accutase (Innovative Cell Technologies) for 5-10 minutes, spun down and aspirated, and rescued in mTeSR Plus with 10μM ROCK inhibitor (Y-27632, Stemgent). 500,000 cells were added to a 10cm dish with 10mL of mTeSR Plus with 10μM Y-27632. After 24 hours, cells were switched to mTeSR Plus without Y27632. Cells were fed with fresh mTeSR Plus every day for optimal growth and expanded for a total of 4 days before experimental seeding.

Lentivirus was grown in Lenti-X 293T HEK cells (Takara Bio). HEK cells were grown in deep 10cm tissue culture dishes (Nunc) and fed with 10% fetal bovine serum (FBS, Millipore) in DMEM/F12 (Thermo). Cells were passaged at a dilution ratio of 1:10 every three days with 0.25% Trypsin-EDTA (Life Technologies) according to manufacturer’s instructions.

Bacterial transformations were performed in NEB Stable Competent *E. coli* (NEB C204OH) according to manufacturer’s instructions.

### Soft lithography and microcontact printing

Photomasks were designed in AutoCAD and submitted to Artnet Pro (formerly CAD/Art) for printing. For the neural tube model, circles were arranged in hexagons with with a diameter of 250 μm per circle and edge-to-edge circle spacing of 100 μm. For transduction efficiency optimization experiments, circles were arranged in 4×4 format with a diameter of 350 μm per circle and inter-circle spacing of 1000 μm. For 24-well format, 24 sets of hexagons or 4×4 grids were arranged with an inter-well spacing of 9 mm for compatibility with 96-well and multichannel pipette spacing.

Microcontact printing was performed as in Anand et al., 2023. Briefly, 50 μm thick 73 mm diameter dry photoresist films (ADEX, DJ Microlaminates) were laminated onto silicon wafers (University Wafers) using a SKY 335R6 laminator (SKY-DSB Co., Ltd) at a temperature of 65°C. Photomasks were placed on top of the photoresist under a long pass filter (PL-360LP, Omega) and then cured under a 365 nm UV lamp (Uvitron) for 16 seconds at of 25 mW/cm^2^ intensity. Silicone and photoresist were developed in cyclohexanone (Sigma) for 5 minutes before rinsing with acetone and then isopropanol, then drying with a stream of compressed air. Resulting molds were baked on a glass dish on a Cimarec hot plate (Thermo) 150°C for 1 hour and then placed in a 10cm Petri dish and silanized with 50 µl of Trichloro(1H,1H,2H,2H-perfluorooctyl)silane (Sigma) on a glass slide in a vacuum chamber overnight. PDMS (Ellsworth Adhesives) was mixed at a curing agent:base reagent ratio of 1:10 and vacuum-treated for 30 to 60 minutes to remove air bubbles. 20g to 40g of PDMS were poured into a Petri dish containing the silicone mold and the dish was baked in an oven (VWR, Sheldon Manufacturing) at 80°C for 2 hours. Stamps were cut out of the dish in a biosafety cabinet in the shape of their rectangular circumference and sterilized by UV treatment under a germicidal lamp for 20 minutes.

Matrigel hESC-qualified Matrix (354277, Corning) was diluted at double the concentration recommended by manufacturer’s instructions in DMEM/F12 media. 5 mL of diluted Matrigel was added to each stamp and left to polymerize overnight at room temperature before seeding.

### 24-well design and construction

24-well sticker templates were designed in Silhouette studio with a well diameter of 8 mm and an inter-well distance of 9 mm, for compatibility with 96-well and multichannel pipette spacing. Stickers were printed using a Silhouette Cameo cutting machine.

PDMS was mixed and vacuum-treated as above. 20g of PDMS was poured into an empty Petri dish and the dish was baked at 80°C for 2 hours. PDMS was cut out from the dish in a biosafety cabinet and placed in the Petri lid for cutting. The 24-well sticker template was applied, and wells were cut out in a biosafety cabinet using an 8 mm biopsy punch (Ted Pella). Well systems were UV-treated for 20 minutes on each side for disinfection.

### Neural tube organoid differentiation

For neural tube organoid protocol optimization, cells were seeded on glass coverslips as in Anand et al., 2023. Briefly, glass coverslips were cut to fit wells of a 6-well plate, sprayed with ethanol, and dried with compressed air in a biosafety cabinet. Matrigel-coated PDMS hexagon stamps were rinsed in distilled water, dried with compressed air, and applied to the center of the coverslip for several minutes in order for Matrigel transfer to occur. Gentle pressure was applied to the stamp to help with transfer. The stamp was then removed for seeding.

Expanded hESCs were rinsed with PBS and treated with Accutase for 15 minutes at room temperature. Cells were then collected, spun down for 3 minutes at 200rcf, aspirated, resuspended in at 3 million cells/mL in mTeSR Plus with 10μM Y27632. The well was filled with cells suspended at 1-2 million cells/mL and incubated for 1 hour in a tissue culture hood to form attachment to the Matrigel. Wells were then rinsed with DPBS (Lonza), the well system was removed, and the plate received one final rinse in DPBS before addition of fresh mTeSR Plus with 10μM Y27632. 24 hours post-seed, media was changed to mTeSR Plus without Y27632.

Neural tube differentiation was initiated at 48 hours post-seed (Day −2) in N2B27 media. Small molecule and signaling protein concentrations were optimized as described in Figure S1B-C. The optimized protocol was as follows: Ectodermal differentiation was initiated at Day −2 with N2B27 + 10 µM LDN + 5 µM SB + 1 µM IWP3 + 5% Matrigel by volume. On Days 0 and 2, media was replaced with N2B27 + 10 ng/mL BMP4 + 5 µM SB + 3 µM IWR1 + 5% Matrigel by volume. Total volume for a 6-well plate was 2 mL, while the total volume for a 10cm plate was 12 mL.

### Plasmid design and cloning

A dual guide lentiviral transfer plasmid backbone was designed according to Replogle et al., 2022, with the payload hU6-guide1-CR3^CS1^-mU6-guide2-CR1^CS2^-EF1alpha-mCerluean. Two guide sequences against each gene were selected according to Replogle et al., 2022. Transfer plasmids were cloned as in Replogle et al., 2022, but modified to utilize arrayed oligo annealing and ligation. Briefly, a backbone containing hU6-CR1^CS2^-EF1alpha-mCerluean was double digested with BsmBI-v2 (NEB) between the hU6 and CR1 and treated with Quick CIP (NEB). Oligo strands were annealed and ligated with backbone and a gBlock containing CR3^CS1^-mU6 using T4 ligase (NEB) as previously described ^57^, but in arrayed 96-well format. NEB Stable (NEB) bacteria were transformed with each plasmid in 96-well plates using heatshock for 30s at 42°C in a thermocycler. Bacteria were rescued in 1 ml SOC media (Thermo Fisher) and grown in 5 ml LB media (Thermo Fisher) with 100 µg/ml carbenicillin overnight at 30°C on a rotating roller drum before miniprep (Qiagen). For clonal plasmid preparation, 50 µl from 5 ml overnight cultures were plated uniformly on on LB + Carbenicillin plates (Teknova) using autoclaved glass beads, grown for 24-48h at 30C, and re-inoculated into overnight liquid culture prior to miniprep. Double digest assay was performed with DraIII-HF (NEB) and BamHI-HF (NEB). Sanger sequencing of the insert was performed using a forward or reverse primer flanking the insert site via Genewiz from Azenta. Whole plasmid sequencing was performed via Plasmidsaurus. We were unable to clone a plasmid with guides against NR6A1 that showed any CFP expression, perhaps due to toxicity of the guides.

### Lentiviral production in 96-well format

Lentiviral particles carrying each gRNA were grown in Lenti-X 293T HEK cells in 96-well tissue culture plates (Celltreat). HEK cells were first passaged via Trypsin according to manufacturer’s instructions and seeded at 60,000 cells/well. HEK cells were transfected when 60% to 95% confluent, about 24 hours post-seed. Transfection reagents were prepared in 96-well PCR plates (VWR). A master mix of 13 ng/μL pMDLg/pRRE, 9 ng/μL pRSV-Rev, and 4.3 ng/μL pMD2.G was deposited into each well at 3 μL/well. pMDLg/pRRE, pRSV-Rev, and pMD2.G were gifts from Didier Trono (Addgene plasmid # 12251; http://n2t.net/addgene:12251; RRID:Addgene_12251; Addgene plasmid 12253; http://n2t.net/addgene:12253; RRID:Addgene_12253; Addgene plasmid 12259; http://n2t.net/addgene:12259; RRID:Addgene_12259). Transfer plasmids at 17.6 ng/μL were deposited into each well at 3 μL/well. A master of jetPrime reagent diluted 1:16 in jetPrime buffer was deposited in each well at 3 μL/well, and all 9 μL of well content were thoroughly mixed and incubated for 10 minutes. After incubation, 91 μL of 10% FBS-DMEM/F12 was added to each reagent well. HEK plates were aspirated, and media was replaced with 100 μL 10% FBS-DMEM/F12 and transfection reagents. Four hours later, BSL2+ safety media change was performed into 70 µL of fresh 5% FBS-DMEM/F12. Virus was allowed to grow for 48 hours before transduction.

### Microcontact printing and hPSC seeding in 24-well format

24-well systems were prepared as follows in a biosafety cabinet immediately before seeding: A 50mm by 75mm borosilicate glass coverslip (Brain Research Laboratories) was sprayed with ethanol, dried with compressed air, and placed in the lid of a deep 10cm tissue culture dish (Nunc). Matrigel-coated stamps were aspirated, rinsed in MilliQ water, dried with compressed air, and applied to the center of the coverslip for several minutes in order for Matrigel transfer to occur. Gentle pressure was applied to the stamp to help with transfer. The 10cm plate bottom was applied to hold the coverslip in place, and the plate was up to mark the corners of the stamp with pen on the bottom the plate. The plate was then returned to the biosafety cabinet, the plate bottom taken off, and the stamp removed carefully to keep the coverslip aligned with the markings. The 24-well PDMS was applied to the coverslip in alignment with the stamp markings and pressed forcefully to create a seal with the coverslip. A diamond-tip scribe (Techni-tool) was used to trim the lateral edges of the coverslip and the well system was then transferred to the plate bottom.

Seeding was performed in a BSL2+ biosafety cabinet. Expanded hESCs were rinsed with PBS and treated with Accutase for 15 minutes at room temperature. Cells were collected, spun down for 3 minutes at 200 x g, aspirated, resuspended in at 3 million cells/mL in mTeSR Plus with 10μM Y27632, and transferred to a pipetting reservoir. Using a multichannel pipette, 50 μL of cell suspension was added to 24 wells of a 96-well plate (CellTreat) for viral mixing. 50 μL of viral supernatant was added to each well and mixed gently before transfer of the full 100 μL to each well of the 24-well system for seeding. Cells were incubated for 1 hour in a tissue culture hood to form attachment to the Matrigel. Wells were then rinsed with DPBS with Ca^2+^/Mg^2+^ (Lonza), the well system was removed, and the plate received one final rinse in DPBS before addition of fresh mTeSR Plus with 10μM Y27632. 24 hours post-seed, media was changed to mTeSR Plus without Y27632.

### Immunostaining

Live organoids were rinsed in DPBS, fixed for 1 hour at room temperature in 4% Formaldehyde, and rinsed three times with PBS. Organoids were fixed between Day 1 (one day of BMP4) treatment and Day 4 (four days of BMP4 treatment) for timecourse analysis. All terminal morphologies were analyzed at Day 4. Because of organoid size, each primary and secondary antibody staining step was done at 4°C on a shaker set to 100 rpm for 48 hours under foil. Organoid morphologies were assayed with Alexa-647-conjugated NCAD antibody (CST 99377S) and Alexa-488-conjugated ECAD antibody (CST 3199). ZO1 was stained with FITC-conjugated primary antibody Invitrogen 33-9111. TFAP2A was stained with primary antibody DSHB 3B5 and secondary antibody Invitrogen A31571. OCT4 was stained with primary antibody CST 2840 and secondary antibody Invitrogen A31573.

### Epifluorescence imaging of fixed and live samples

Epifluorescence microscopy was performed on a Zeiss AxioObserver Z1 inverted microscope with Zeiss 10x and 20x NA 1.3 plan apo objectives. Images were acquired with an Orca-Flash 4.0 CMOS camera (Hamamatsu). LED light was filtered using Zeiss filter sets 43 HE DsRed, 46 HE YFP, 47 HE CFP, 49 DAPI, and 50 Cy5.

For live imaging, a live imaging lid with a CO2 nozzle was constructed for a 10cm plate. Cells were imaged with 37°C incubation and 5% CO2. Transduction efficiency was examined at 48 hours post-seed with live epifluorescence microscopy in phase and 47 HE CFP channels. For live timelapses of actin, organoids were incubated in media containing 50 nM SiR-actin for 24 hours, then imaged for the indicated timeframes after BMP4 addition.

### Confocal imaging of fixed samples

Confocal microscopy was performed on a Zeiss LSM 980 with Airyscan using a Zeiss 10x NA 0.45 objective. Z-stacks were performed through whole organoids. Single slices through the middle of each organoid are presented in figures.

### Single-cell RNA sequencing library preparation

Neural tube organoids were collected at Days 2 and 4 (after 48 hours or 96 hours of BMP4 treatment, respectively) for dissociation. Day 4 organoids perturbed with different guides were pooled for sequencing. Organoids were scraped off glass coverslips into a new 10cm dish, diluted in 5mL of PBS, collected, and spun down for 2 minutes at 200 x g. PBS was then aspirated until there was 0.5mL left in the tube, and cells were resuspended and transferred with a wide bore pipette into a fresh 10cm dish. Cells were then incubated in Accutase in a tissue culture hood for 20 minutes, then pipetted up and down with a narrow bore pipette to break up clumps. Single cells were spun down for 3 minutes at 200 x g, aspirated, and resuspended at 1.1 million cells/mL in PBS + 0.04% BSA.

Cells were counted on the LUNA-FX7 Automated Cell Counter (Logos Biosystems) using fluorescence detection for viability with an acridine orange/propidium iodide stain (Part No. F23011). All samples had viability greater than 85%, with concentration ranges from 600 to 1500 cells/µL. After counting, all samples were loaded into Chip G per the user guide from 10x Genomics (Part No. CG000316). GEMs were formed targeting 3,000 to 22,000 cells and reverse transcription completed immediately after. The *Feature cDNA Primers 3* (Part No. 2000289) were used to amplify cDNA from poly-adenylated mRNA and sgRNA containing either capture sequence 1 or capture sequence 2. Size selection was used to separate the amplified cDNA molecules for 3’ Gene Expression and for CRISPR screening library construction. The amplified cDNA was verified via TapeStation (4200 TapeStation instrument, Agilent Technologies) using High Sensitivity D5000 tape and reagents (Part No. 5067-5592 & 5067-5593).

The amplified cDNA from poly-adenylated mRNA was fragmented, end repaired, and A-tailed followed by adaptor ligation, and PCR amplification with each sample receiving a unique set of dual indices (Part No. 1000215). The cDNA from sgRNA molecules was further amplified and the CRISPR Screening libraries were then generated by PCR amplification with each sample receiving a unique set of dual indices (Part No. PN-1000242).

Final libraries were diluted and ran using the High Sensitivity D5000 tape and reagents (Part No. 5067-5592 & 5067-5593) on the 4200 TapeStation (Agilent Technologies). Libraries were quantified via Kapa qPCR using the Complete Universal Kit (Part No. 07960140001, Roche Sequencing Solutions) and the CFX96 Touch Real-Time PCR Detection System (Bio-Rad Laboratories). Libraries were sequenced on an Illumina NovaSeq 6000 instrument using the parameters outlined in the user guide (Read1: 28 bp, i7 index: 10 bp, i5 index: 10 bp, Read2: 90 bp). After sequencing and demultiplexing data are analyzed with Cell Ranger Count pipeline.

### Quantification of transduction efficiency

Transduction efficiency analysis was performed on live epifluorescence microscopy of micropatterned stem cells, 48 hours after transduction and immediately before 3D cyst induction with Matrigel. Ilastik was trained to recognize the circular outline of each micropatterned colony in the phase channel and to return the center coordinates and diameters (d) of each colony. Circle parameters were then applied to the mCerulean channel in Python to extract mCerulean pixel values both from the colony and from a 2D background halo (d+10 to d+35 pixels). Colony pixels were categorized as mCerulean+ if they reached expression above the 95th percentile value in the background halo. Some micropatterns were located under the CO_2_ adapter during live imaging and thus their transduction efficiency could not be measured.

For nuclear OCT4 knockdown analysis, nuclear centers were selected via DAPI stain in Fiji. Ten-pixel positions surrounding and including the nuclear center were extracted in Python from the OCT4 channel and summed to score OCT4 intensity. OCT4 values of each nucleus (n>600 per micropatterned circle) were binned into histograms. An OCT4-nucleus was defined as having lower than the 5^th^ percentile of nuclear OCT4 values in CFP-nuclei.

### Organoid morphology analysis

Organoid morphology analysis was performed on epifluorescence microscopy images of Day 4 organoids using the Ilastik software platform. User-generated labels and epifluorescence were used to distinguish neural tube tissue (NCAD-positive) and non-neural tissue (ECAD-positive) through machine-learning pattern-based pixel classification. Morphological characteristics were extracted from the segmented objects, including area measurements. For organoid size quantification, adjacent NCAD- and ECAD-positive regions were summed to determine total organoid area. For ECAD/NCAD area, the former areas were divided.

### Single-cell RNA-sequencing analysis of Day 2 and Day 4 organoids and Week 3 and 4 human embryos

After sequencing and demultiplexing, FASTQ files were run through the 10X Cell Ranger Count pipeline to align reads to the GRCh38-2020 reference genome, make guide calls according to capture sequencing, and produce a cell by gene count matrix.

For Day 2 and 4 organoids, Week 3 embryos, and Week 4 embryos ^49^, gene expression was analyzed in Python with Scanpy (v1.10.2). For each dataset, cells with too few or too many reads (potential doublets) were excluded from analysis based on manual cutoffs. All other cells were analyzed with sparse multimodal decomposition (SMD, Melton & Ramanathan, 2021) to find informative genes for clustering. Cells were then clustered using Scanpy’s Louvain or Leiden algorithm in the space of the top 100 to 250 SMD genes (excluding ribosomal, mitochondrial, and cell cycle factors) with a clustering resolution of 0.2-0.5. In the case of Week 3 and Week 4 embryos, multiple rounds of clustering were performed to first identify non-ectodermal cell types and re-cluster ectodermal cell types only. In all cases, clusters were identified based on expression of canonical markers, including transcription factors. Cells were assigned to neural plate, neural plate border, and surface ectoderm for subsequent analysis. Raw reads of included cells were converted to transcripts per million (TPM) and log1p-normalized prior to further analysis.

Mediolateral axis analysis of Day 2 data was performed on log1p-normalized reads with Scanpy. First, principal component coordinates for each cell were calculated with scanpy.tl.pca using SciPy’s ARPACK solver. Cells were color-coded along principal components 1 and 2 by their Louvain cluster identifications to find the neural-to-non-neural orientation along principal component 1. A nearest neighbors’ distance matrix was then generated using scanpy.pp.neighbors with a local neighborhood size of 20. Diffusion pseudospace coordinates for each cell were calculated with scanpy.tl.dpt, using the least-neural cell on the principal component axis as the root cell.

For Day 4 organoids, guide calls and counts were merged with the cell by gene count matrix according to cell barcode in Python. Guide conditions were selected for analysis if at least ten cells were sequenced from that guide condition (12 out of 18 non-scramble guides met this criteria). Knockdown efficacy of each guide condition was measured against a scramble guide control condition using an Anderson Darling test (SciPy) with a significance cutoff of p<0.05. Gene expression of neural genes in each selected guide condition was compared to scramble control. Genes were defined as differentially expressed if they passed an Anderson Darling with p<0.05.

### Motif enrichment analysis

Gene names that were strongly downregulated in *ZIC2*-perturbed organoids (log2(mean fold change relative to scramble control) <-0.04) were bulk-converted to ENSEMBL IDs using g.Profiler. Genomic regions (10kb upstream of start site and 100bp regions downstream of start site) were extracted using the Eukaryotic Promoter Database (EPD). Regions were analyzed with MEME-SEA for enrichment of transcription factor motifs relative to scrambled sequences. Enrichment scores of transcription factor families with multiple enriched motifs were combined for a final family enrichment score.

## Notes

https://figshare.com/articles/dataset/2026_HuangAnand/31151509

https://github.com/royahuang/2026_HuangAnand

### Summary of Updates

This manuscript has been updated to address reviewer comments. Figures 1 and S1 have been updated to include more thorough analyses of timepoints from in vivo human neural data. The Figure 1 caption has been updated to specify geometry of organoids and our interpretation of lumen stain data. Two supplementary videos have been added showing shedding of dead cells into the lumen. The Methods section has been updated to include details of mediolateral axis analysis from scRNA-seq data.

